# A self-limiting, TREM2-dependent anabolic program confers microglial resilience in Alzheimer’s disease

**DOI:** 10.64898/2026.09.16.752098

**Authors:** Da Lin, Jeffrey R. Atkinson, Weidong Wu, Min Chen, Sohan Jayasekara, Benjamin M. Segal, Anjun Ma, Jie Gao

## Abstract

Microglia are key drivers of Alzheimer’s disease (AD), and TREM2, one of the strongest genetic risk factors, enables the disease-associated microglia (DAM) state required for plaque engagement. How TREM2 sustains this protection, and why it falters as disease advances, remain unclear. Using proteomics, single-cell transcriptomics, and in vivo metabolic labeling, we show that plaque-associated microglia mount a TREM2-dependent anabolic program coupling nascent protein synthesis to mitochondrial biogenesis. This program, not the DAM signature, marks phagocytically competent microglia; its loss is associated with proteostatic overload and mitochondrial dysfunction. Unexpectedly, this anabolic program is self-limiting, peaking at low amyloid burden and declining as burden rises. This decline is conserved in humans: across 523 donors, anabolic capacity tracks TREM2 but falls at advanced Braak stage even as DAM signature keeps rising. Thus, anabolic capacity, rather than activation state, marks microglial resilience in AD, and sustained stimulation of phagocytosis by anti-amyloid antibodies risks biosynthetic exhaustion and self-limiting efficacy.

## Introduction

In Alzheimer’s disease (AD), microglia serve as a primary line of defense, tasked with the containment and clearance of neurotoxic Aβ aggregates ^1^. Genetic studies have placed microglial function at the center of AD risk, implying that a robust innate immune response helps delay pathogenesis ^2^ ^3^. This protective capacity, however, is not static. As pathology progresses, microglia undergo profound phenotypic changes, initially mounting a containment response, but frequently transitioning toward dysregulated, senescent, or exhausted states that fail to limit neurodegeneration ^4^ ^5^. Understanding the mechanisms that allow microglia to maintain functional resilience under chronic proteotoxic stress is therefore a priority for therapeutic intervention.

Recent advances in single-cell transcriptomics have mapped the trajectory of this response, defining a conserved ‘Disease-Associated Microglia’ (DAM) or ‘Microglial Neurodegenerative Phenotype’ (MGnD) signature ^6^ ^7^. This program, driven largely by TREM2 (triggering receptor expressed on myeloid cells 2), involves downregulation of homeostatic checkpoints (e.g., *P2ry12*, *Cx3cr1*) and induction of lipid-sensing and phagocytic pathways (e.g., *Apoe*, *Lpl*, *Clec7a*). Loss-of-function variants (e.g., R47H) underscore the importance of TREM2, markedly increasing AD risk and are associated with impaired microglial clustering around plaques and diffuse, neurotoxic amyloid pathology ^8^ ^9^ ^10^ ^11^. Although the DAM/MGnD framework has been highly influential, human studies describe a broader spectrum of microglial states, indicating that transcriptional state alone may not fully capture functional capacity. Indeed, the acquisition of a transcriptional signature is only a blueprint: executing the functions of plaque-associated microglia (proliferation, migration, and continuous phagocytosis) imposes a formidable biosynthetic and bioenergetic demand, requiring constant synthesis of new membranes, hydrolytic enzymes, and organelles to replace those consumed during Aβ degradation. How microglia acquire the metabolic resources to sustain this state in the AD brain remains poorly understood. While studies have well characterized immunometabolism in peripheral macrophages, where activation couples to distinct metabolic switches (e.g., glycolysis vs. oxidative phosphorylation) ^12^, the metabolic dependencies of plaque-associated microglia are far less explored. Although TREM2 has been linked to microglial metabolic fitness ^13^, whether it supports a spatially organized anabolic program in vivo, and whether such a program is separable from the DAM transcriptional state, remains unknown.

Here, we combine integrated proteomics, transcriptomics, single-cell RNA sequencing, and in vivo metabolic labeling to characterize the biosynthetic and bioenergetic features of microglial resilience to Aβ pathology. We find that Aβ pathology is accompanied by a TREM2-dependent anabolic program, a coordinated increase in protein synthesis and mitochondrial biogenesis that we resolve in situ at the phagocytic interface, and that this program, rather than the DAM transcriptional signature, distinguishes phagocytically competent microglia. Unexpectedly, the program is self-limiting: it peaks in microglia with low amyloid burden and is progressively suppressed as burden rises, in both mouse and human tissue. In the absence of TREM2, microglia do not sustain this response and instead exhibit proteostatic overload and mitochondrial dysfunction. We integrate these observations into a supply-demand model of microglial clearance and consider its implications for anti-amyloid immunotherapy, whose efficacy depends on the same phagocytic capacity.

## Results

### Microglia mount a TREM2-dependent anabolic program in response to Aβ pathology

To gain mechanistic insight into microglial adaptation to amyloid pathology, we performed integrated proteomic and transcriptomic profiling of primary microglia isolated from *App^NL-G-F^* and *App^NL-G-F^*;*Trem2*^KO^ mice at 3 and 9 months of age, corresponding to early and intermediate stages of amyloid deposition, respectively (Fig. 1A).

**Figure 1:**
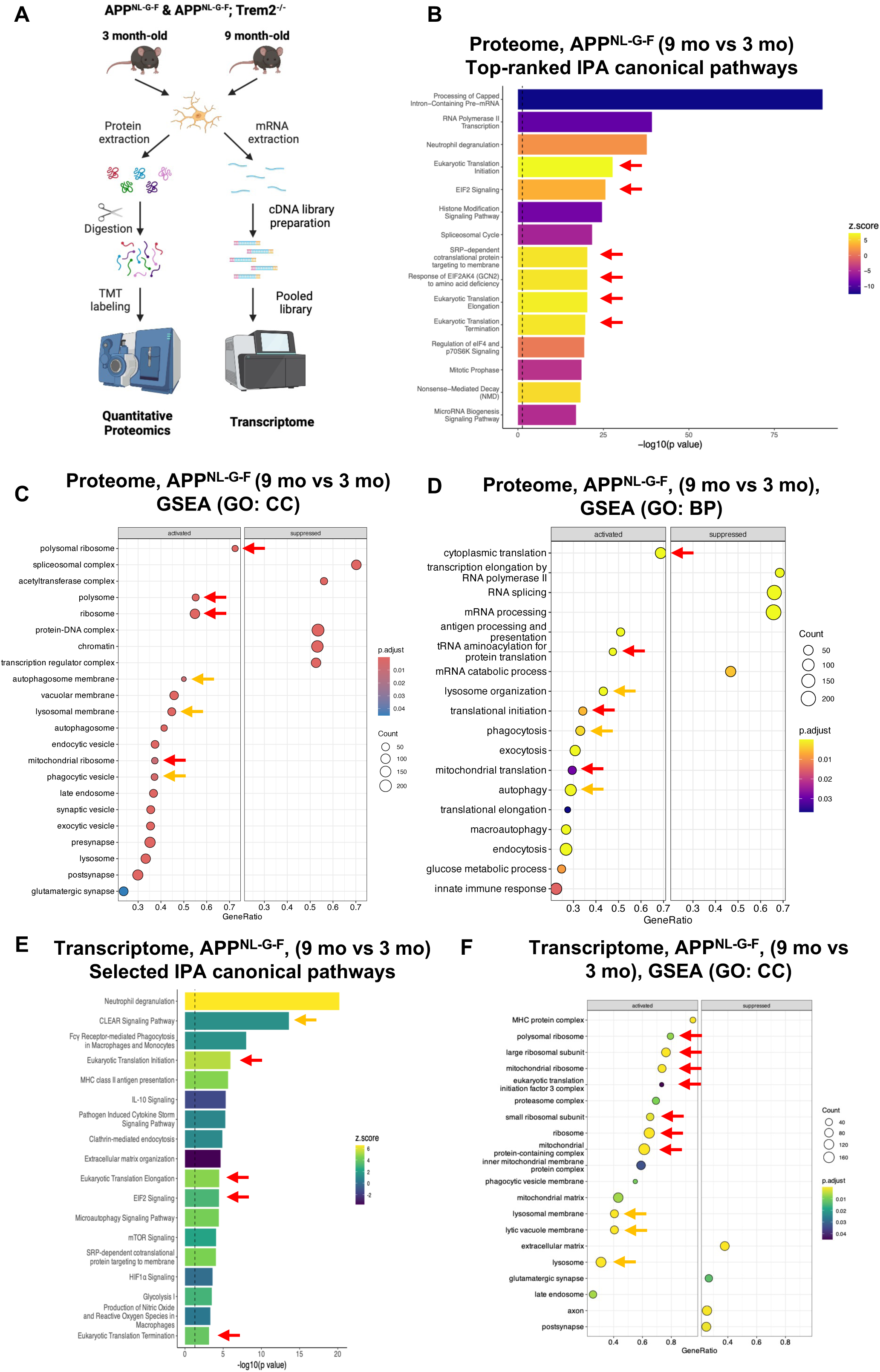
Microglia mount a proteome-wide anabolic adaptation to Aβ pathology. **(A)** Experimental schematic. Primary microglia were isolated from 3-and 9-month-old *App^NL-G-F^*and *App^NL-G-F^;Trem2^KO^* mice for integrated proteomic and transcriptomic profiling. **(B)** Ingenuity Pathway Analysis (IPA) of differentially abundant proteins (DAP) in 9-month-old *App^NL-G-F^* vs. 3-month-old *App^NL-G-F^* microglia, highlighting the enrichment of protein-synthesis pathways (red arrows). **(C–D)** Gene Set Enrichment Analysis (GSEA) dot plots for Cellular Component (C) and Biological Process (D) terms. Red arrows indicate the concurrent enrichment of “Ribosome” (anabolic) signature and orange arrows indicate “Lysosomal membrane” (catabolic) signatures enriched in 9-month-old vs. 3-month-old *App^NL-G-F^* microglia. **(E)** IPA and **(F)** GSEA of differentially expressed gene (DEG) in 9-month-old *App^NL-G-F^*, highlighting genes involved in Protein Translation (red arrows) and catabolic processes (orange arrows).

In *App^NL-G-F^* microglia, proteomic profiling identified over 7,000 proteins, of which 2,621 were differentially abundant between 3 and 9 months, indicating extensive proteome remodeling as pathology progressed. Ingenuity Pathway Analysis (IPA) revealed a robust induction of protein synthesis pathways, including Eukaryotic Translation Initiation, Elongation, and Termination, and EIF2 Signaling (Fig. 1B, red arrows). Gene Set Enrichment Analysis (GSEA) of both Cellular Component (CC) and Biological Process (BP) terms confirmed the increased abundance of translation machinery, including cytosolic and mitochondrial ribosomal subunits (Fig. 1C-D, red arrows; Extended Fig. 1A). In parallel, proteins associated with degradative processes, such as ‘autophagosome membrane’, ‘lysosomal membrane’, and ‘phagocytic vesicles’, were also enriched (Fig. 1C–D, orange arrows). Transcriptomic profiling corroborated this adaptive response, albeit more modestly, revealing increased expression of genes associated with protein translation (Fig. 1E, red arrows), ribosome biogenesis and mitochondrial respiratory-chain complexes (Fig. 1F; Extended Fig. 1B, red arrows), and lysosomal/CLEAR pathways (Fig. 1E–F, orange arrows). Together, these data indicate that microglia undergo coordinated metabolic remodeling during Aβ pathology, coupling elevated protein synthesis with enhanced degradative capacity to sustain proteome renewal.

To determine whether this metabolic remodeling is TREM2-dependent, we compared the profiles of *App^NL-G-F^* and *App^NL-G-F^*;*Trem2*^KO^ microglia. The impact of TREM2 was tightly linked to disease stage. At 3 months, when Aβ burden is minimal, TREM2 deficiency produced no significant change in the microglial proteome (Extended Fig. 2A). By 9 months, however, loss of TREM2 profoundly disrupted the proteome, resulting in 1,480 significantly altered proteins (Fig. 2A). In both proteome and transcriptome, loss of TREM2 markedly reduced the disease-associated microglia (DAM) signature, such as Gpnmb, Cst7, and Lpl (Fig. 2A, B), consistent with the critical role of TREM2 in driving the conversion of homeostatic microglia into DAM under Aβ stress.

**Figure 2:**
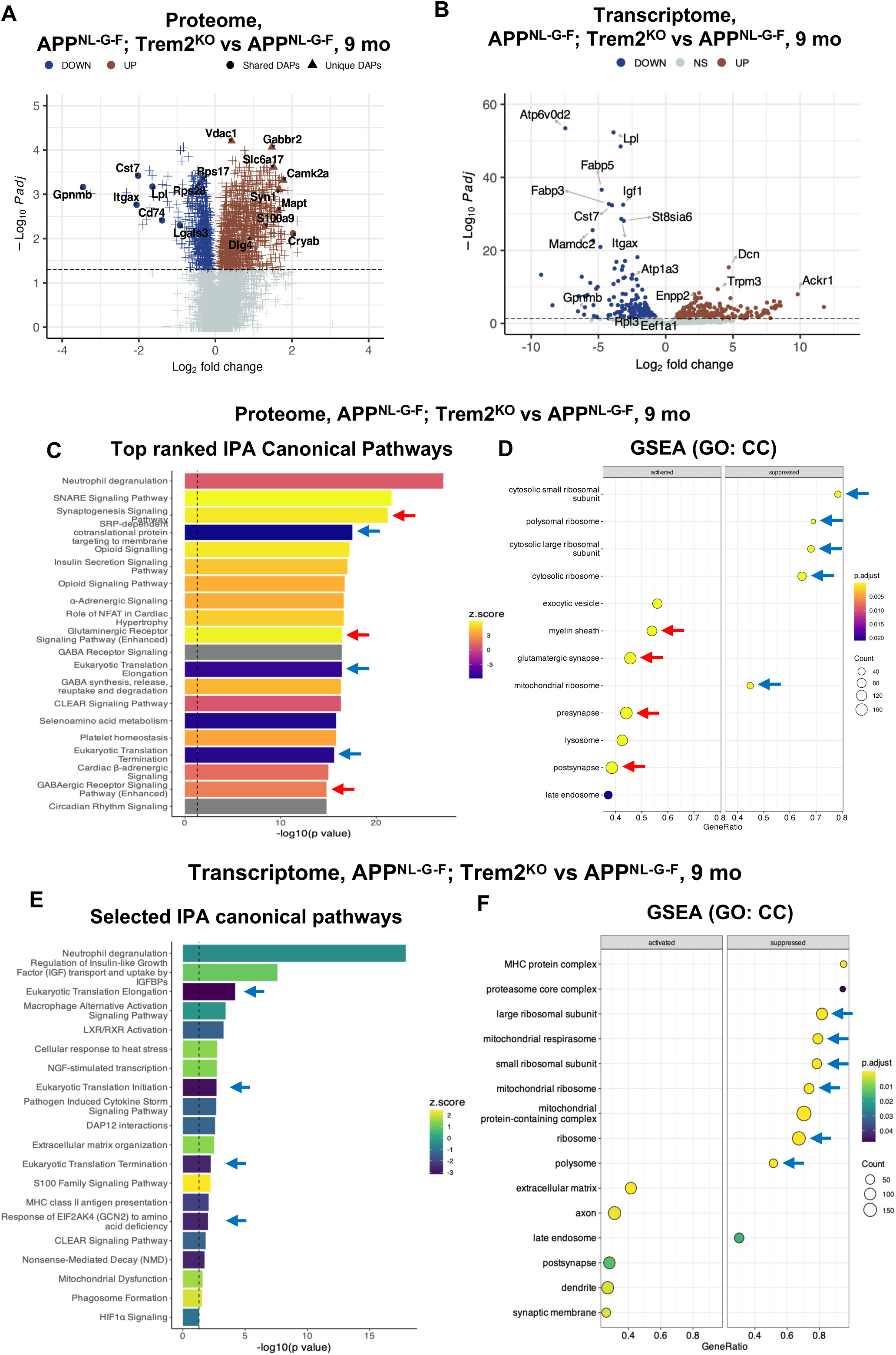
TREM2 deficiency causes a collapse of the anabolic response. **(A)** Volcano plot comparing the proteome of 9-month-old *App^NL-G-F^* and *App^NL-G-F^;Trem2^KO^*microglia. Triangle dots indicate uniquely altered proteins in the *App^NL-G-F^;Trem2^KO^* microglia. The Venn diagram shows the unique and shared genes and proteins between the transcriptome and proteome. **(B)** Volcano plot comparing the transcriptome of 9-month-old *App^NL-G-F^* and *App^NL-G-F^;Trem2^KO^* microglia. **(C)** IPA of top-ranked pathways in *App*^NL-G-F^;*Trem2*^KO^ microglia, showing specific suppression of protein translation pathways (blue arrows) and the increased levels of synaptic proteins (red arrows). **(D)** GSEA dot plots confirming the broad reduction of cytosolic and mitochondrial ribosomal components in *Trem2^KO^* microglia (blue arrows); red arrows indicate the concurrent accumulation of synaptic and myelin signatures. **(E)** IPA and **(F)** GSEA of differentially expressed gene (DEG) in 9-month-old *App^NL-G-F^*and *App^NL-G-F^;Trem2^KO^* microglia, highlighting suppressed genes involved in protein translation and ribosome (blue arrows).

Beyond the DAM signature, a dominant effect of TREM2 deficiency was the collapse of the anabolic response. Proteome IPA revealed marked downregulation of translation-related pathways, including SRP-dependent protein targeting, translation elongation, and translation termination (Fig. 2C, blue arrows). GSEA confirmed broad reductions in cytosolic and mitochondrial ribosomal components, as well as ribosome biogenesis (Fig. 2D; Extended Fig. 2B, blue arrows), indicating that TREM2 is required for the induction of these anabolic pathways during Aβ pathology. Transcriptomic profiling reinforced these findings, showing significantly reduced expression of genes involved in protein translation, ribosomal assembly, and mitochondrial respiratory complexes (Fig. 2E-F; Extended Fig. 2C). Directional integration of proteomic and transcriptomic datasets revealed that, while App^NL-G-F^ microglia coordinate protein translation, transport, and degradation, Trem2^KO^ microglia fail to mount this proteome remodeling response (Extended Fig. 2D-E). Consistent with defective proteome renewal, *Trem2*^KO^ microglia accumulated synaptic and myelin proteins, materials normally cleared following phagocytosis (Fig. 2C, D, red arrows). This proteostatic failure was evident in the proteome, where many of the uniquely upregulated DAPs were synaptic proteins (Fig. 2A) and proteome IPA and GSEA enriched synaptic and myelin terms. Together, these findings indicate that microglia adapt to Aβ pathology through a TREM2-dependent anabolic program that is associated with proteome remodeling and proteostatic maintenance.

### TREM2 drives nascent protein synthesis in plaque-associated microglia

To validate the anabolic program revealed by our omics analysis in vivo, we employed Fluorescent Non-Canonical Amino Acid Tagging (FUNCAT) to label newly synthesized proteins (Fig. 3A). Sixteen hours after intraperitoneal injection of azidohomoalanine (AHA), a methionine analog incorporated into nascent proteins, we analyzed CX3CR1+ CD45+ microglia by flow cytometry (Extended Fig. 3A). At 3 months of age, FUNCAT intensity was comparable between *App^NL-G-F^* and *App^NL-G-F^*;*Trem2*^KO^ microglia. By 9 months, however, microglia from *App^NL-G-F^* mice exhibited an approximately twofold increase in protein synthesis, whereas TREM2 deficiency abolished this induction (Fig. 3B-C). Histograms revealed a distinct ‘FUNCAT-high’ shoulder in *App^NL-G-F^*microglia (Fig. 3B, red arrow), indicating the emergence of a metabolically active subpopulation.

**Figure 3:**
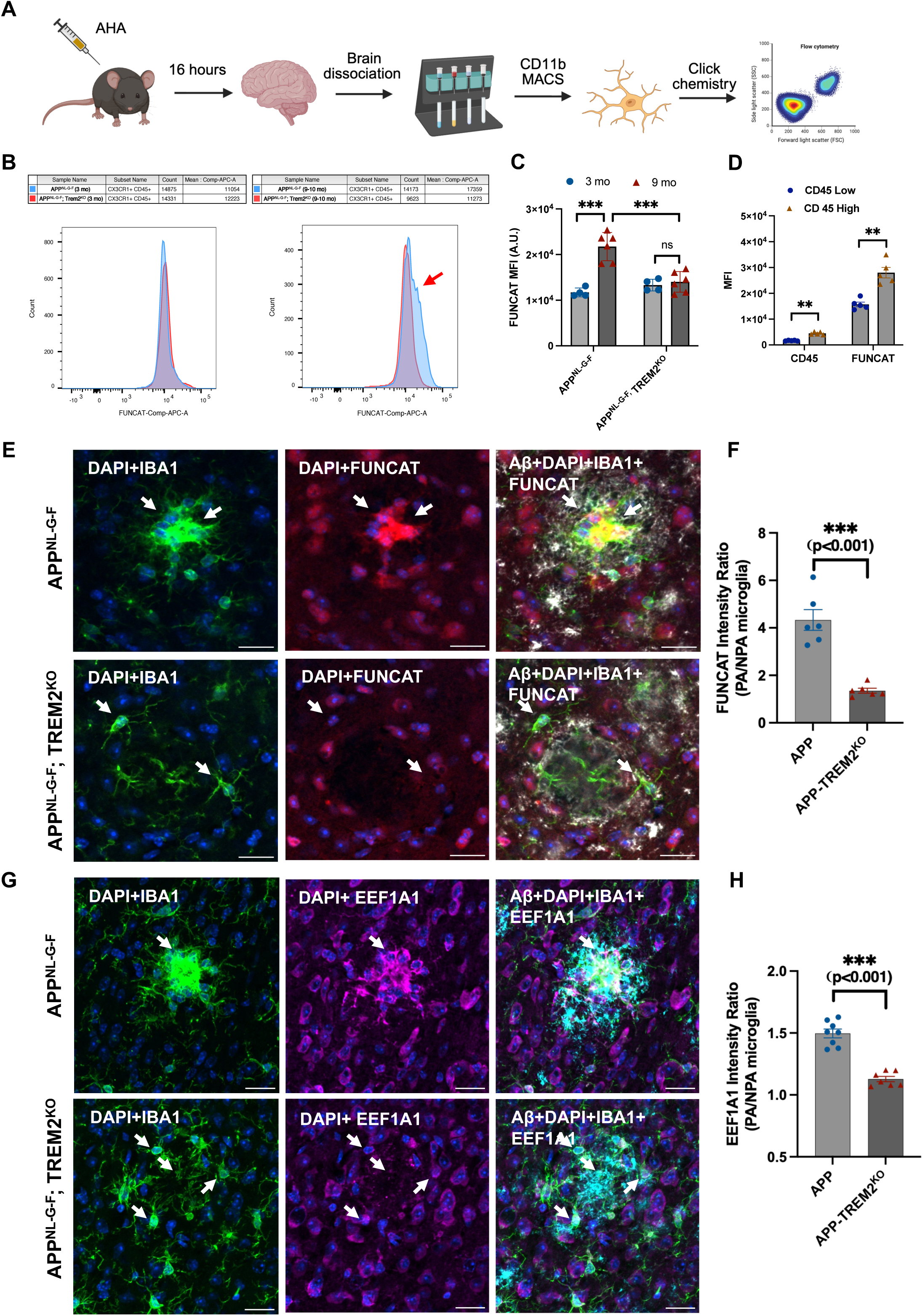
TREM2 drives nascent protein synthesis in plaque-associated microglia. **(A)** Schematic of *in vivo* FUNCAT labeling. Mice were injected with Azidohomoalanine (AHA, 50 mg/kg) 16 hours prior to microglia isolation and analysis. **(B)** Representative flow cytometry histograms of FUNCAT intensity in CX3CR1^+^ CD45^+^ microglia. Red arrow indicates the “FUNCAT-high” shoulder present in *App^NL-G-F^* microglia but absent in *Trem2^KO^*mice. **(C)** Quantification of mean fluorescence intensity (MFI) for FUNCAT signal. **(D)** Comparison of CD45 and FUNCAT intensity between CD45^high^ and CD45^low^ microglial populations. **(E)** Representative confocal images of cortical plaques stained for Iba1 (green), Aβ (82E1, grey), and nascent proteins (FUNCAT, red). Scale bar: 50 µm. **(F)** Quantification of FUNCAT intensity in plaque-associated microglia. **(G)** Representative confocal images of the translation elongation factor EEF1A1 (Magenta) in plaque-associated microglia from 9-month-old *App^NL-G-F^* and *App^NL-G-F^;Trem2^KO^* mice, co-stained for Iba1 (green) and Aβ (Cyan). White arrows indicate EEF1A1 induction within Iba1+ plaque-associated microglia. Scale bar: 20 µm. **(H)** Quantification of EEF1A1 intensity within Iba1+ plaque-associated microglia. n = 7-8 mice per group. ***P < 0.001 by two-tailed unpaired t-test.

Because CD45^high^ microglia are known to accumulate around plaques ^14^, we examined their relationship to protein synthesis. While the CD45^high^ subpopulation expanded with the progression of Aβ pathology (Extended Fig. 3B-C), this expansion was largely absent in *Trem2*KO mice. CD45^high^ microglia showed markedly higher FUNCAT intensity than CD45^low^ microglia (Fig. 3D), suggesting that plaque-associated microglia are the primary population undergoing elevated nascent protein synthesis. In situ FUNCAT labeling confirmed a robust induction of protein synthesis specifically in the plaque-associated microglia of *App^NL-G-F^* mice, an induction largely abolished in *App^NL-G-F^*;*Trem2*^KO^ mice (Fig. 3E-F). To corroborate this induction with a direct component of the translational machinery, we stained for eukaryotic translation elongation factor 1α1 (EEF1A1), one of the anabolic-signature genes. EEF1A1 was robustly induced within plaque-associated microglia of *App^NL-G-F^*mice but not in those of *App^NL-G-F^*;*Trem2*^KO^ mice (Fig. 3G, H), mirroring the FUNCAT results and directly linking the anabolic program to elevated translational capacity in situ. These results are consistent with the previously established TREM2-mTOR axis ^13^: phosphorylated ribosomal protein S6 (p-RPS6), a canonical marker of mTORC1 activity, was likewise increased in plaque-associated microglia in a TREM2-dependent manner (Extended Fig. 3D, E). Together, these findings indicate that the TREM2-dependent anabolic program is accompanied by increased translational capacity and nascent protein synthesis in plaque-associated microglia.

### The anabolic adaptation is TREM2-dependent and self-limiting across the Aβ load gradient

Having established that plaque-associated microglia mount anabolic adaptation, we asked how it scales with phagocytic burden. We profiled the transcriptomes of Aβ-engaged microglia using Methoxy-X04 (X04), a fluorescent probe that selectively labels fibrillar Aβ (Fig. 4A). Primary microglia from *App^NL-G-F^* and *App^NL-G-F^*;*Trem2*^KO^ mice were sorted into X04⁺ (Aβ-phagocytic) and X04⁻ (non-phagocytic) populations. To capture the full spectrum of Aβ load, X04⁺ microglia were further stratified into low-, medium-, and high-intensity groups (Fig. 4B). At early stages of pathology (5-6 months), the proportion of X04⁺ microglia was comparable between genotypes, indicating that TREM2 is not required for the initial engulfment of Aβ fibrils. By the mid-stage (9-10 months), however, TREM2 deficiency markedly reduced the total X04⁺ population, driven largely by the specific loss of the highly phagocytic X04^high^ subset (Fig. 4B-C). Because X04 intensity reflects cumulative Aβ uptake and processing, these findings indicate that TREM2 deficiency either impairs the survival of the high-Aβ-load (X04^high^) subpopulation or alters Aβ uptake and intracellular processing kinetics, under chronic Aβ burden.

**Figure 4:**
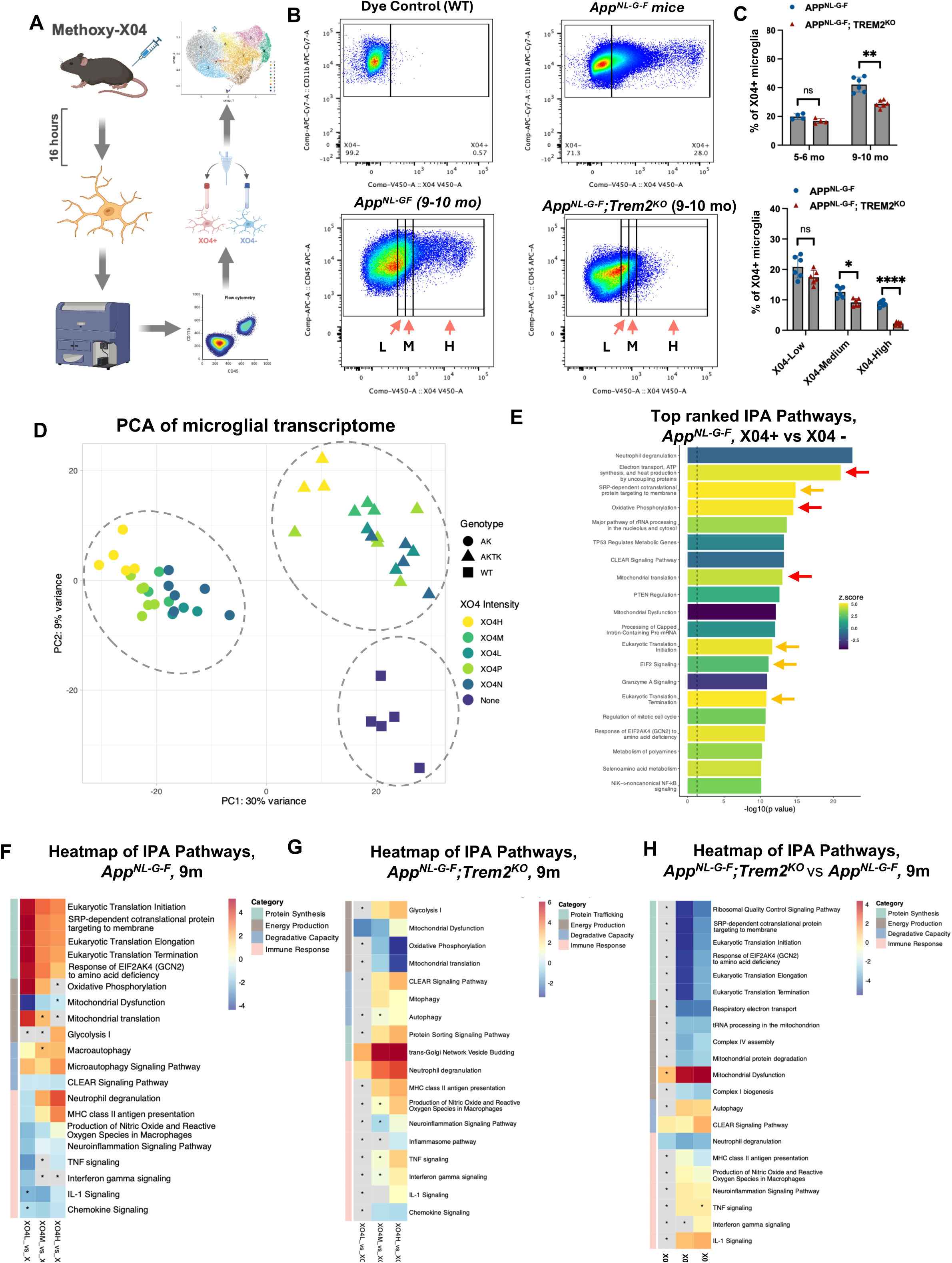
The microglial anabolic adaptation is TREM2-dependent and self-limiting across the Aβ load gradient. **(A)** Schematic of the gating strategy used to sort Methoxy-X04 (X04) positive (X04^+^) and negative (X04^-^) microglia for transcriptional profiling. **(B)** Representative flow cytometry plots illustrating the gating of Aβ-phagocytic microglia. Upper panels: Comparison of Wild-Type (WT) and *App^NL-G-F^* mice confirms the specificity of the X04 signal. Lower panels: Stratification of the X04^+^ population into Low, Medium, and High fluorescence intensity bins, representing increasing phagocytic load, in *App^NL-G-F^* and *App^NL-G-F^;Trem2^KO^* mice. **(C)** Temporal analysis of Aβ-phagocytic microglia. Upper panels: Expansion of the X04^+^ population in *App^NL-G-F^* mice from 5–6 months to 9–10 months of age. Lower panels: Comparison of X04 intensity distributions in 9-month-old *App^NL-G-F^* and *App^NL-G-F^;Trem2^KO^*mice, showing the marked reduction of X04^high^ population in TREM2^KO^ microglia. **(D)** Principal Component Analysis (PCA) of bulk RNA-seq samples (*n* = 46). Dots represent individual biological replicates across genotypes: WT (*n* = 5), *App^NL-G-F^*(*n* = 23), and *App^NL-G-F^;Trem2^KO^* (*n* = 18). Colors denote X04 classification (X04^-^ vs. X04^+^ subsets: Low, Medium, High). Note that X04 levels drive separation along PC2, defining distinct plaque-phagocytic states. **(E)** IPA of top-ranked pathways enriched in X04^+^ vs. X04^-^ *App^NL-G-F^*microglia. Arrows highlight the robust induction of Mitochondrial Biogenesis/Oxidative Phosphorylation (red) and Protein Targeting/Synthesis (orange) upon Aβ engagement. **(F, G)** IPA Comparison Analysis illustrating transcriptional changes of microglia across the phagocytic gradient (X04^high^, X04^med^, X04^low^) relative to X04^-^ controls in *App^NL-G-F^* mice (F) and at *App^NL-G-F^;Trem2^KO^* mice (G) at 9 months of age. **(H)** IPA Comparison of *App^NL-G-F^*vs *App^NL-G-F^;Trem2^KO^* stratified by Methoxy-X04 intensity at 9 months of age. The activation z-scores for the selected significantly enriched canonical pathways (Fisher’s exact test; adjusted *P* < 0.05) are presented as a heatmap. Color scale indicates predicted pathway directionality: red corresponds to positive z-scores (predicted activation), blue represents negative z-scores (predicted inhibition), and grey denotes pathways for which IPA was unable to predict directionality. Pathways lacking statistical significance are marked with an asterisk (*). Rows of pathways are grouped into four categories according to their biological relevance.

Transcriptomic profiling of X04⁺ and X04⁻ microglia from 9-month-old wild-type, *App^NL-G-F^*, and *App^NL-G-F^*;*Trem2*^KO^ mice revealed clear genotype-specific clustering by principal-component analysis (PCA) (Fig. 4D). Within each genotype, samples also aligned along the X04-intensity gradient, indicating a shift of transcriptional states proportional to Aβ load (Fig. 4D; Extended Fig. 4A-B). Differential expression analysis between X04⁺ and X04⁻ microglia in *App^NL-G-F^* mice revealed a strong induction of mitochondrial pathways, including Electron Transport Chain, Oxidative Phosphorylation, and Mitochondrial Translation (Fig. 4E, red arrows), as well as protein translational programs, such as SRP-dependent protein targeting, Translation Initiation, and Translation Termination (Fig. 4E, orange arrows). These results indicate that Aβ phagocytosis is accompanied by a coordinated anabolic program that couples protein synthesis with mitochondrial biogenesis. Loss of TREM2 abolished this transcriptional program (Extended Fig. 4C). Critically, analysis across the X04 subpopulations (X04⁻, low, medium, high) revealed that the program is non-monotonic: induction of anabolic genes, including those for protein synthesis and mitochondrial function, peaked in X04^low^ microglia and declined progressively as phagocytic burden increased (Fig. 4F). In contrast, pathways associated with the DAM signature, such as Neutrophil Degranulation and MHC II Antigen Presentation, increased with X04 intensity, while classical inflammatory pathways (TNF, IFN-γ) were suppressed or unchanged (Fig. 4F), indicating that anabolic activation reflects a metabolically adaptive, largely non-inflammatory response. Loss of TREM2 abolished the anabolic response in X04^low^ and X04^medium^ microglia (Fig. 4G); as Aβ load increased, *Trem2*^KO^ microglia displayed reduced expression of genes related to oxidative phosphorylation and mitochondrial translation, increased signatures of mitochondrial dysfunction, a compensatory upregulation of glycolysis, and heightened pro-inflammatory signaling (Fig. 4G). Across the X04 gradient, TREM2 loss had minimal effects on non-phagocytic microglia but led to a marked reduction of the anabolic program, including genes related to protein synthesis and mitochondrial bioenergetics, in the phagocytic X04^low^ and X04^medium^ populations (Fig. 4H). These results define the anabolic program as TREM2-dependent and self-limiting: it is engaged upon Aβ phagocytosis, peaks in microglia carrying a low amyloid burden, and is progressively suppressed as burden rises, so that the biosynthetic capacity that sustains clearance is lowest in the most heavily Aβ engaged cells. TREM2 loss prevented the program from being sustained even at low burden and precipitated metabolic failure under chronic Aβ stress.

### Anabolic capacity, better than DAM signature, marks phagocytically competent microglia

To place anabolic adaptation within the broader landscape of microglial activation, we performed single-cell RNA sequencing on X04⁺ (phagocytic) and X04⁻ (non-phagocytic) microglia isolated from 9-10-month-old *App^NL-G-F^* mice. Clustering identified the expected spectrum of transcriptional states ^15^ ^16^, including Homeostatic (HM), Activated Response (ARM), and Transition Response (TRM) microglia (Extended Fig. 5A-B). However, projecting the canonical Disease-Associated Microglia (DAM) signature onto this map revealed that DAM genes were expressed broadly across both X04⁺ and X04⁻ populations (Extended Fig. 5C). This indicates that acquisition of the transcriptional DAM signature is insufficient to distinguish microglia functionally engaged in phagocytosis from those merely responsive to the pathological environment.

To resolve the molecular determinants of phagocytic competence, we next examined transcriptional heterogeneity within these clusters. UMAP visualization revealed that X04⁺ and X04⁻ microglia occupied distinct regions within the nominally defined HM and TRM clusters (Extended Fig. 5D). Increasing the clustering resolution resolved this heterogeneity, splitting HM into subclusters 0 (non-phagocytic-enriched) and 2 (phagocytic-enriched), and TRM into subclusters 6 (non-phagocytic) and 1 (phagocytic) (Fig. 5A, B). DAM Module Scores were similar between phagocytic (Clusters 1, 2) and non-phagocytic (Clusters 6, 0) pairs (Extended Fig. 5E), indicating that the DAM signature correlates poorly with phagocytic state. IPA and GSEA of DEG between Clusters 1, 2 and Clusters 6, 0 revealed that the phagocytic state was defined by a synchronized upregulation of genes involved in ribosome biogenesis, protein synthesis, and mitochondrial oxidative phosphorylation (Fig. 5C; Extended Fig. 5F, G, H), mirroring the proteomic anabolic signature identified earlier. The signature of anabolic adaptation features the induction of the translation elongation factor Eef1a1 and structural ribosomal subunits (*Rpl10*, *Rpl13*), coupled with respiratory chain subunits spanning the input (*Ndufa1*), catalytic core (*Cox4i1*), and ATP-generating output (*Atp5e*) of the electron transport chain (Fig. 5D). These anabolic signatures were higher in the phagocytic-enriched microglia (subclusters 1 and 2) and distinct from the DAM signature. This anabolic induction appeared distinct from inflammatory activation: neuroinflammatory signaling was reduced in the phagocytic clusters (1, 2) relative to their non-phagocytic counterparts (6, 0) (Extended Fig. 5F, G).

**Figure 5:**
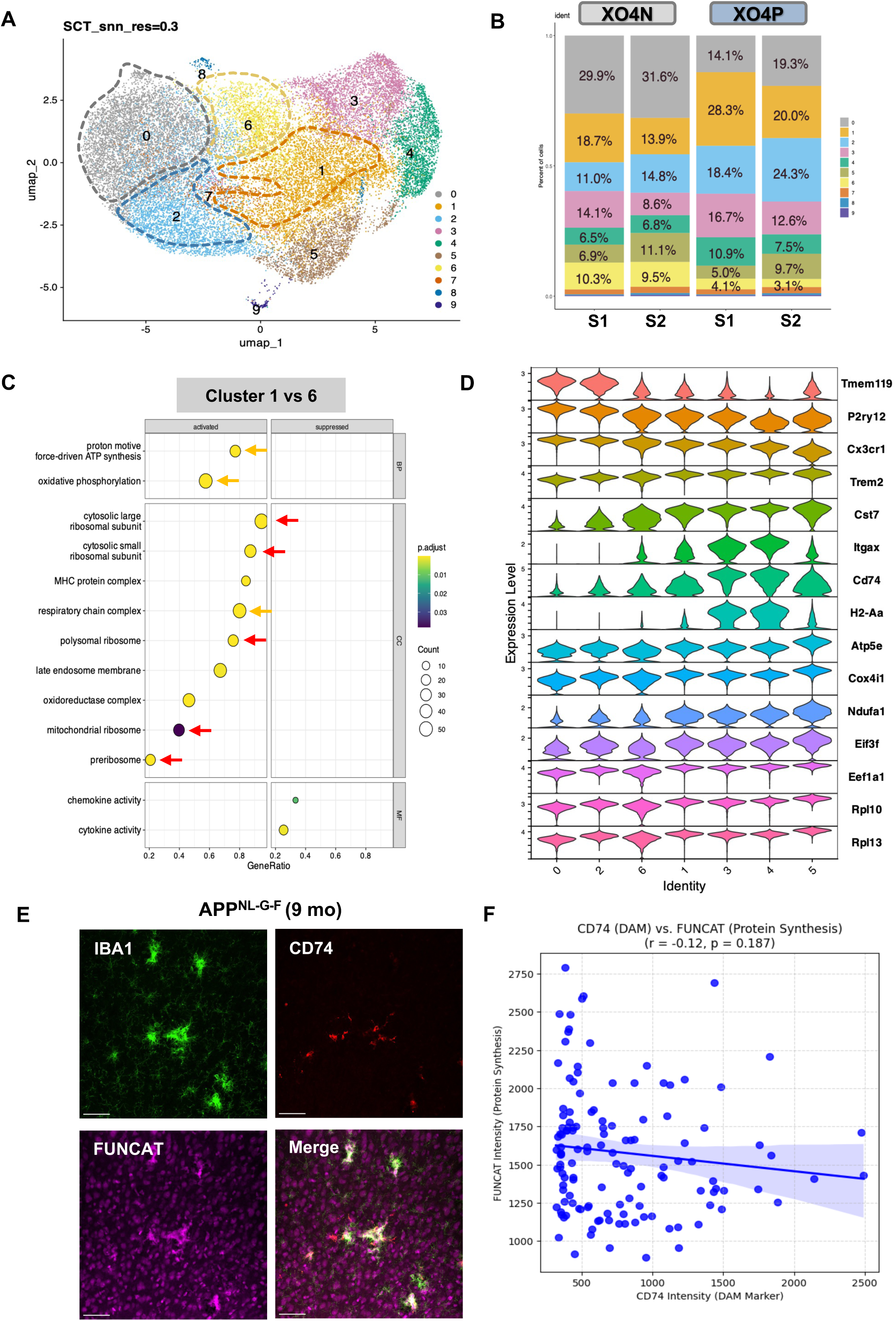
The Anabolic Adaptation signature defines phagocytic competence more accurately than the canonical DAM signature. **(A)** UMAP projection of single-cell RNA-seq profiles (resolution 0.3) from sorted X04+ (phagocytic) and X04-(non-phagocytic) microglia isolated from 9-10 month-old *App^NL-G-F^*mice. High-resolution subclustering splits the canonically defined Homeostatic (HM) and Transition (TRM) clusters into phagocytic-and non-phagocytic-enriched subpopulations (HM into clusters 0 and 2; TRM into clusters 1 and 6) that segregate by phagocytic competence. **(B)** Quantification of cluster composition, showing the relative enrichment of each subcluster within the X04+ versus X04-fractions. **(C)** Gene Set Enrichment Analysis (GSEA) comparing the phagocytic-enriched subcluster (Cluster 1) versus the non-phagocytic subcluster (Cluster 6), revealing robust enrichment of Ribosome and Oxidative Phosphorylation pathways in the phagocytic subpopulation. **(D)** Violin plots showing expression of the Homeostatic signature (Tmem119, P2ry12, Cx3cr1), DAM signature (Trem2, Cst7, Itgax, Cd74, H2-Aa), and Anabolic Adaptation signature (Atp5e, Cox4i1, Ndufa1, Eif3f, Eef1a1, Rpl10, Rpl13) across the identified subpopulations. While the DAM signature is comparable between phagocytic and non-phagocytic states, the Anabolic Adaptation signature is selectively upregulated in the phagocytic clusters (1 and 2). **(E)** Representative confocal images of plaque-associated microglia in 9-month-old *App^NL-G-F^*mice co-stained for Iba1, the DAM marker CD74, and FUNCAT (nascent protein synthesis). Most Iba1^high^ microglia show robust FUNCAT signal, whereas CD74 is induced only in a subset. Scale bar: 50 µm. **(F)** Quantification of the correlation between FUNCAT and CD74 intensity within individual Iba1+ microglia (Pearson’s R =-0.12, P = 0.187), showing that the transcriptional DAM state does not predict anabolic activity.

Finally, we spatially validated the relationship between the DAM signature and anabolic activity in situ. We co-stained brain sections from 9-month-old *App^NL-G-F^*mice for Iba1, the canonical DAM marker CD74, and the nascent protein label FUNCAT. While most plaque-associated (Iba1^high^) microglia exhibited robust anabolic activity (high FUNCAT signal), CD74 was induced only in a subset of plaque-associated microglia (Fig. 5E), and CD74 intensity showed no correlation with FUNCAT signal within Iba1+ microglia (Pearson’s R = −0.12, P = 0.187; Fig. 5F). Collectively, these data indicate that the anabolic adaptation signature, rather than the DAM signature, better distinguishes the phagocytic microglial state, separating the functional capacity to engage with stress from the broader transcriptional response to Aβ.

### Loss of the anabolic program is associated with proteostatic overload

Our omics results suggest that the ability to fuel phagocytosis defines microglial competence; if so, its loss should manifest as a failure of phagocytic processing. Consistent with this notion, and with proteome data showing the synaptic and myelin proteins that accumulated in the *Trem2*^KO^ microglia (Fig. 2; Extended Fig. 6A), we examined the intracellular accumulation of phagocytic cargo directly. Confocal imaging of the postsynaptic marker PSD95, together with Iba1 and Aβ, revealed a substantial buildup of PSD95 puncta within *Trem2*^KO^ microglia near plaques (Fig. 6A, B). These inclusions were surrounded by CD68+ lysosomes (Extended Fig. 6B), indicating stalled degradation of engulfed synaptic material. The presynaptic marker vGluT1 similarly accumulated in *Trem2*^KO^ microglia (Extended Fig. 6C, D). This defective clearance extended beyond synaptic debris: MBP staining showed a striking increase in intracellular myelin accumulation in *Trem2*^KO^ microglia at plaque sites (Fig. 6C, D). Because PSD95 and MBP are not amyloid-derived, their accumulation is unlikely to be explained solely by differences in Aβ uptake between genotypes. The simultaneous buildup of synaptic and myelin proteins, despite the presence of lysosomal markers, suggests a general failure of phagocytic degradation when TREM2-dependent anabolic support is absent.

**Figure 6:**
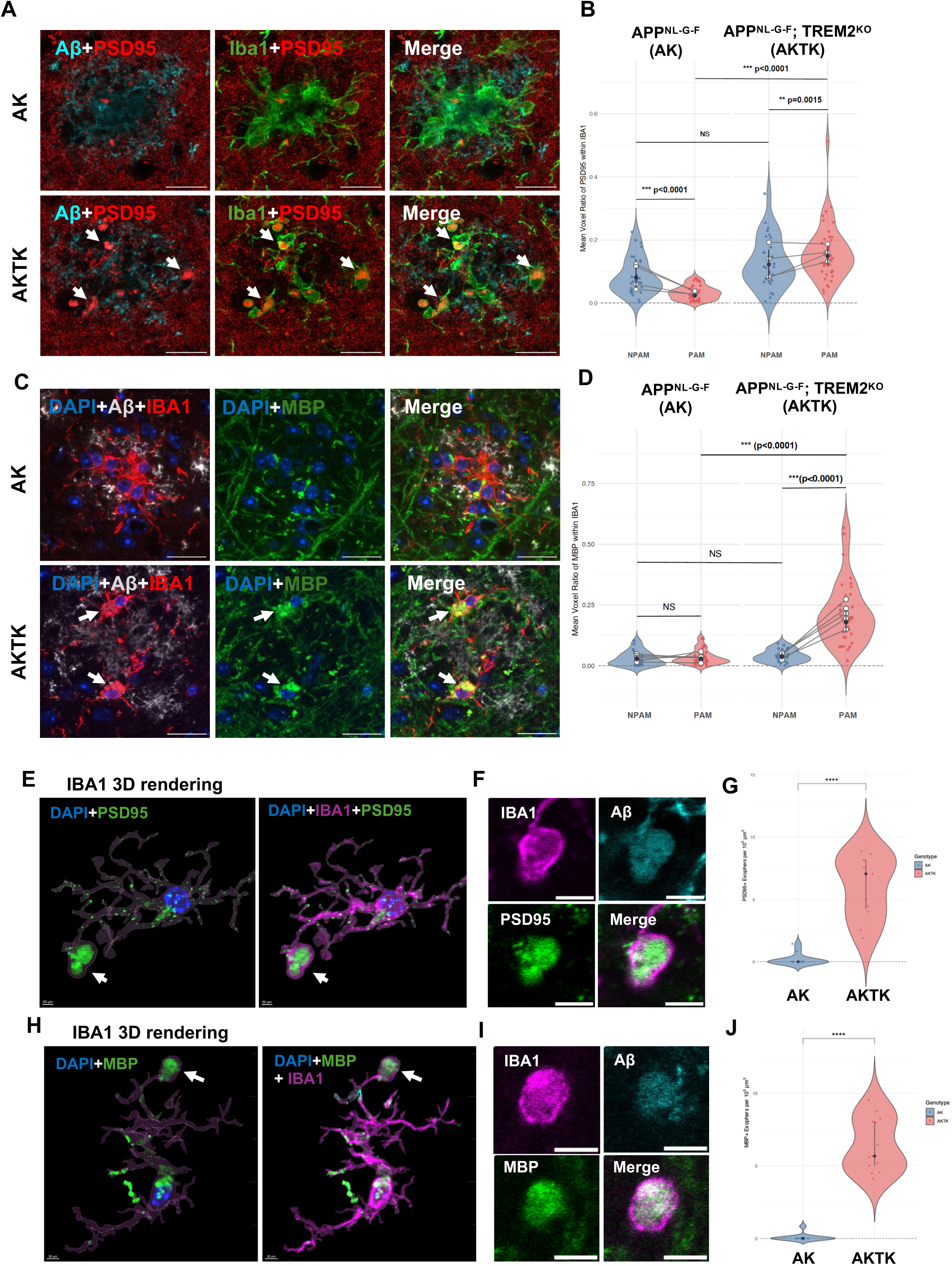
Loss of TREM2 leads to the intracellular accumulation of undigested cargo. (A, **C)** Representative confocal images of plaque-associated microglia in 9-month-old mice co-stained for PSD95 (postsynaptic marker, A) or Myelin Basic Protein (MBP, C), together with Iba1 and Aβ. White arrows highlight the pronounced intracellular accumulation of PSD95 and MBP puncta within the soma of *Trem2^KO^* microglia. Scale bar: 50 μm. **(B, D)** Quantification of cargo accumulation relative to plaque proximity. Violin plots display the voxel overlap percentage of PSD95 (B) or MBP (D) within Iba1+ ROIs, classified as Non-Plaque-Associated (NPAM) or Plaque-Associated (PAM). Colored points represent individual ROIs; solid black circles and error bars represent Estimated Marginal Means (EMMs) ± 95% CIs derived from a beta-binomial Generalized Linear Mixed Model (GLMM). Lines connect paired regions within biological replicates (n = 5-6 mice per group). Pairwise comparisons were performed using the Delta method. Significance levels: *P < 0.05, **P < 0.01, ***P < 0.001, ****P < 0.0001. Exact p-values are shown for non-significant trends. **(E)** 3D surface rendering (Imaris) of a plaque-associated microglial cell in *App^NL-G-F^;Trem2^KO^* mice, co-stained for DAPI, PSD95 (green), Iba1 (magenta), and Aβ (cyan). Note the extrusion of a large, cargo-filled process. **(F)** High-magnification (63X) single optical slice (0.17 μm) confirming the enclosure of PSD95 puncta and Aβ aggregates within the Iba1+ process. **(G)** Quantification of PSD95+ processes in the plaque niche (10^5^ μm^3^ ROI). Each point represents an individual plaque ROI. Medians and interquartile ranges (IQRs) are indicated by dots and bars, respectively. Statistical significance was determined using a non-parametric Wilcoxon rank-sum test with exact permutation (R package coin, v1.4.3). **(H)** 3D surface rendering of processes containing Myelin Basic Protein (MBP) debris. **(I)** Single optical slice (0.17 μm) confirmation of MBP cargo within the process lumen. **(J)** Quantification of MBP+ processes density between genotypes.

In a subset of plaque-associated *Trem2*^KO^ microglia, we additionally observed large (2-8 µm), membrane-bound spherical Iba1^+^/DAPI^-^ vesicles that were filled with PSD95 or MBP (Extended Fig. 6E, F). High-resolution 3D reconstruction from multiple confocal image datasets revealed that these vesicles mostly emerged directly from microglial processes, appearing as segmental enlargements connected by thin membrane nanotubes (Fig. 6E, H; Extended Fig. 6G, H). These enlarged processes were packed with Aβ, along with synaptic (PSD95^+^) and myelin (MBP^+^) debris (Fig. 6F, I), consistent with compromised phagocytic processing and proteostatic overload in TREM2^KO^ microglia. Quantitative analysis revealed a striking increase in the frequency of these events in TREM2^KO^ microglia. Within a standardized plaque-associated volume (10^5^ um^3^), we detected an average of 7-8 PSD95^+^ or MBP^+^ enlarged processes in *App^NL-G-F^*;*Trem2*^KO^ mice, compared to less than 0.5 in *App^NL-G-F^* controls (Fig. 6G, J). When normalized to the number of plaque-associated microglia, we found that 60-70% of *Trem2*^KO^ microglia in the plaque niche were associated with this feature of proteostatic overload, compared to less than 2% of WT microglia, demonstrating a general proteostatic failure upon phagocytic stress in the absence of TREM2.

### Loss of the anabolic program is associated with mitochondrial dysfunction

To determine whether the anabolic defect associated with TREM2 deficiency extends to bioenergetics in vivo, we assessed mitochondrial dynamics using flow cytometry. We reasoned that if TREM2^KO^ microglia fail to mount the anabolic program supporting mitochondrial biogenesis, they would be less able to maintain a healthy organelle pool under the phagocytic stress of amyloid pathology. Freshly isolated microglia were stained with MitoTracker Green (total mitochondrial mass) and MitoTracker Red CMXRos (membrane potential). In *App^NL-G-F^*mice, the CD45^high^ (plaque-associated) population exhibited a coordinated elevation of both mitochondrial membrane potential and lysosomal mass, consistent with the metabolic demands of Aβ clearance (Fig. 7A, B). Interestingly, *Trem2*^KO^ microglia displayed bioenergetic uncoupling: while total mitochondrial mass was significantly increased relative to controls, these organelles exhibited significantly reduced membrane potential. Consequently, the Mitochondrial Health Index (a derived ratio of respiratory-active mitochondria to total mass) was markedly lower in *Trem2*^KO^ microglia (Fig. 7C), indicating an accumulation of metabolically incompetent organelles. Lysosomal mass (LysoTracker) remained unchanged between genotypes (Fig. 7D), suggesting that the accumulation of depolarized mitochondria in *Trem2*^KO^ microglia is unlikely to stem from lysosomal deficiency.

**Figure 7:**
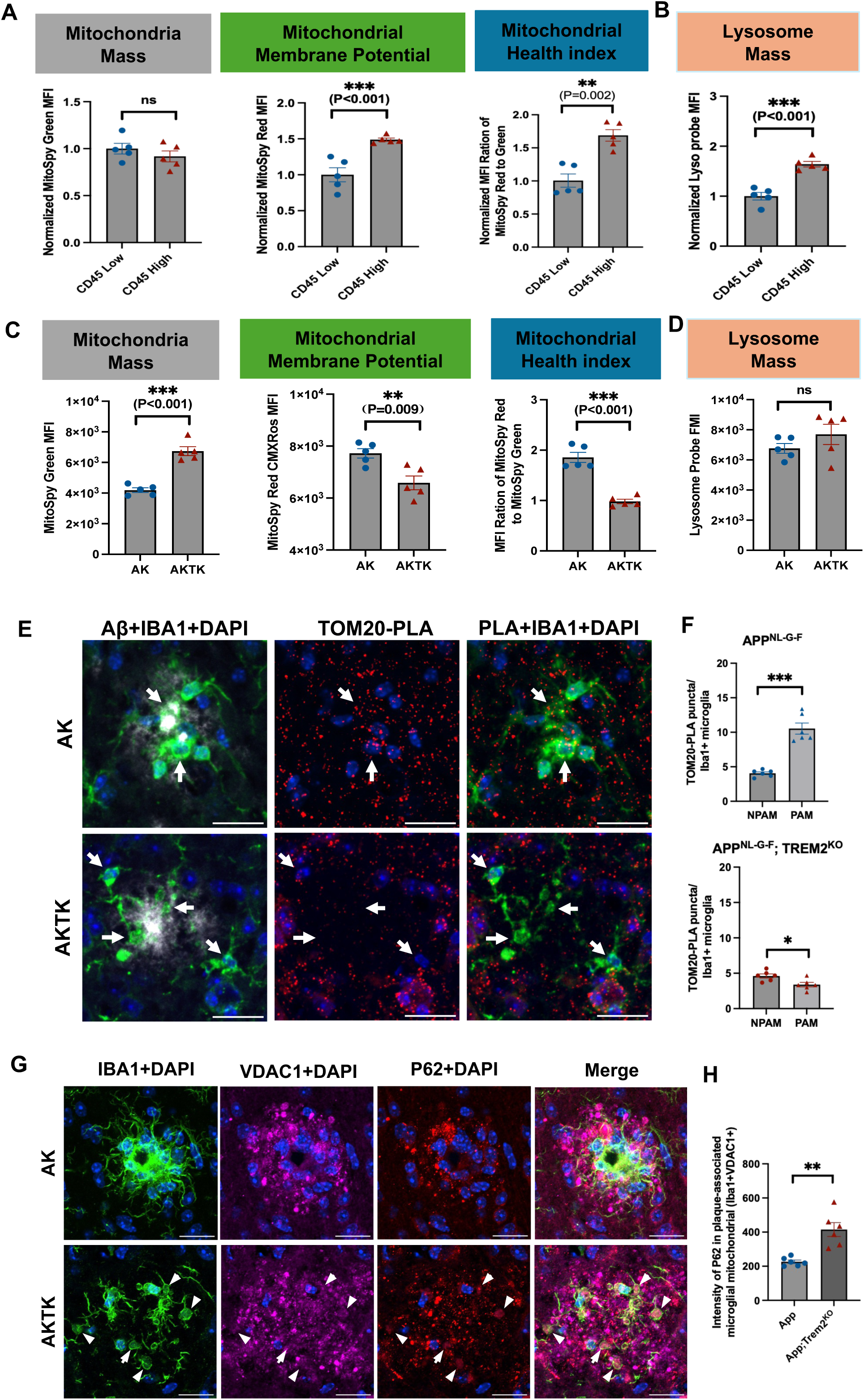
Loss of TREM2 leads to reduced mitochondrial biogenesis and renewal in plaque-associated microglia. **(A)** Flow cytometry analysis of mitochondrial fitness in CD45^low^ (non-plaque) versus CD45^high^ (plaque-associated) microglial from 9-month-old *App^NL-G-F^* mice, quantifying Mitochondrial Mass (MitoTracker Green), Membrane Potential (MitoTracker Red CMXRos), and the Mitochondrial Health Index (ratio of Membrane Potential to Mass). CD45^high^ microglia show increased membrane potential, indicating expanded bioenergetic capacity. **(B)** Lysosomal mass (LysoTracker Deep Red) in the same populations; CD45^high^ microglia show higher lysosomal mass, confirming induction of degradative capacity. **(C)** Mitochondrial fitness in primary microglia from 9-month-old *App^NL-G-F^* and *App^NL-G-F^;Trem2^KO^* mice (same metrics as in A). Trem2^KO^ microglia show a “High Mass / Low Potential” phenotype, indicating accumulation of metabolically incompetent organelles. **(D)** Lysosomal mass (LysoTracker Deep Red); the absence of a genotype difference indicates the mitochondrial accumulation is not driven by lysosomal deficiency. **(E)** Nascent mitochondrial biogenesis in situ by AHA-TOM20 Proximity Ligation Assay (PLA). Representative confocal images show newly synthesized TOM20 (TOM20-PLA puncta, red) within Iba1+ microglia (green) in the plaque niche, robust in *App^NL-G-F^*and blunted in Trem2^KO^ microglia. **(F)** Quantification of nascent TOM20-PLA puncta number per microglia (10-15 randomly selected plaques per mice were quantified). **(G)** Representative confocal images (MIP, 5 μm) of plaque-associated microglia co-stained for VDAC1, P62 (autophagy receptor), and Iba1. White arrows indicate VDAC1+ mitochondria with high p62 in Trem2KO microglia. Scale bar: 20 μm. **(H)** Quantification of P62 intensity in IBA1⁺VDAC1⁺ regions; 20-30 plaques quantified per mouse.

We next asked whether this defect reflects a failure in mitochondrial renewal, particularly in the plaque-associated microglia. To assess biogenesis, we combined metabolic labeling with a proximity ligation assay (PLA). Mice were injected with AHA to tag nascent proteins, followed by click-mediated biotinylation and detection of newly synthesized TOM20 using an anti-biotin/anti-TOM20 PLA antibody pair (Negative control in Extended Fig. 7A). In *App^NL-G-F^* mice, plaque-associated microglia displayed a significant increase in nascent mitochondrial signal (TOM20-PLA+) specifically within IBA1+ regions near plaques (Fig. 7E, F), indicating biogenesis in response to amyloid deposition. In contrast, *Trem2*^KO^ microglia did not increase TOM20-PLA+ signal, indicating that TREM2 is required for the induction of mitochondrial biogenesis. Co-staining of FUNCAT and TOM20 revealed colocalization of nascent protein signal with mitochondria within microglial processes (Extended Fig. 7B), coinciding with increased TREM2 expression in the processes of plaque-associated microglia (Extended Fig. 7C). Despite the reduction in new biogenesis, quantification of confocal images revealed that *Trem2*^KO^ microglia around plaques exhibited increased total mitochondrial mass (% of TOM20+ voxels in Iba1+ area; Extended Fig. 7D, E), corroborating the flow cytometry data and indicating an accumulation of mitochondria that are not being renewed. Co-staining of the outer mitochondrial membrane protein VDAC1 with the autophagy receptor p62/SQSTM1 revealed accumulation of P62⁺/VDAC1⁺ aggregates in plaque-associated TREM2^KO^ microglia (Fig. 7G, H), indicating damaged mitochondria that failed to undergo mitophagy.

### A conserved anabolic program declines in human microglia with AD progression

Our mouse data indicated that a TREM2-dependent anabolic program, distinct from the canonical DAM signature, marks phagocytically competent microglia and is self-limiting. To determine whether this program is relevant to human disease, we examined microglia in an integrated single-nucleus transcriptomic atlas of the human brain (ssKIND) ^17^, restricting the analysis to 523 donors with annotated Braak staging and age (Patient Summary in Table 1). We scored each microglial nucleus for the mouse-derived anabolic module, the canonical DAM signature, and a homeostatic module. Projected onto the integrated microglial manifold, the anabolic, DAM, and homeostatic modules each marked a distinct region of the map, indicating that high-anabolic microglia occupy a transcriptional state separable from both DAM-high and homeostatic-high cells (Fig. 8A). Unsupervised re-clustering resolved a ribosomal/anabolic-high subcluster distinct from a DAM-marker-high subcluster (Extended Fig. 8A), confirming the anabolic program as a distinct transcriptional state in human microglia. To test whether anabolic capacity changes with disease progression, we aggregated microglia to donor-level pseudobulk and compared the anabolic module score across Braak tiers (0–II, III–IV, V–VI), adjusting for donor age. The anabolic score declined with advancing Braak stage (age-adjusted β = −0.41, P = 1.8×10^-5^; Spearman R = −0.35), with the steepest reduction between Braak III-IV and V-VI (Fig. 8B). This decline was not attributable to technical variation: a control module of stably expressed reference genes showed no comparable Braak-dependent change (Extended Fig. 8B), and the median number of genes detected per nucleus remained stable across stages (Extended Fig. 8C). To distinguish loss of a high-anabolic population from a uniform per-cell decrement, we labeled microglia anabolic-high using a global threshold (top 30% of the cell-level anabolic score) and quantified the per-donor anabolic-high fraction across Braak tiers. This fraction contracted sharply between Braak III-IV and V-VI (donor medians 66.7% for Braak III-IV vs. 21.4% for Braak V-VI; P < 1×10^-4^, Spearman R = −0.32; Fig. 8C), whereas the mean score within anabolic-high cells declined only modestly (Fig. 8D); a shift-share decomposition attributed ≈62% of the III–IV→V–VI decline to the reduced fraction of anabolic-high cells and ≈38% to a per-cell decrement (Extended Fig. 8D). Thus the decline is predominantly compositional, reflecting a contraction of the high-anabolic population, with a smaller per-cell component, paralleling the stage-dependent collapse of the anabolic program observed in the mouse.

**Figure 8:**
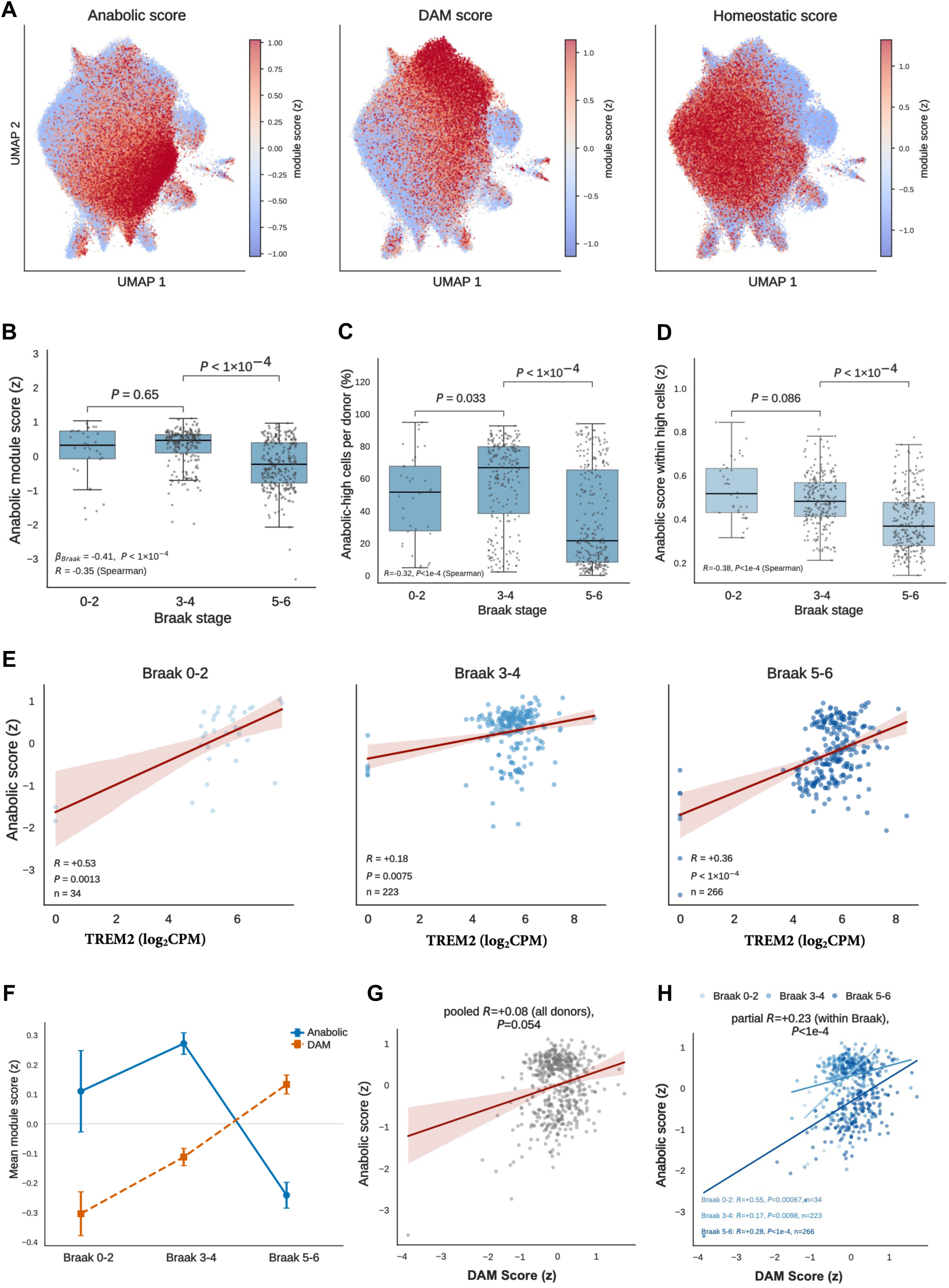
The microglial anabolic program is conserved in humans and declines with AD progression. **(A)** Per-nucleus module scores for the anabolic, DAM, and homeostatic programs projected onto the integrated human microglial UMAP (ssKIND atlas; 523 donors). **(B)** Donor-level pseudobulk anabolic module score across Braak tiers (0–II, III–IV, V–VI). Each point is one donor; boxes show median and interquartile range. The anabolic score declines with Braak stage (linear model β = −0.41, P = 1.8×10⁻⁵; Spearman R = −0.35), most steeply between Braak III–IV and V–VI. n = 34 / 223 / 266 donors (Braak 0–II / III–IV / V–VI). Pairwise comparisons, Mann–Whitney U. (**C**) fraction of anabolic-high cells per donor; (**D**) mean anabolic score within those high cells; cells were labelled anabolic-high at a global threshold (top 30% of the cell-level anabolic score across all cells). **(E)** *TREM2* expression (log₂ CPM) versus anabolic module score across donors, per Braak stage. Spearman R is positive at every stage; partial R controlling for Braak stage = +0.28. Red line, linear fit; shaded band, 95% confidence interval. **(F)** Mean anabolic and DAM module scores across Braak tiers (error bars, s.e.m.), showing divergence by Braak V–VI. Partial Spearman correlations controlling for Braak stage: anabolic– *TREM2* R = +0.28; anabolic–DAM R = +0.23; anabolic–homeostatic R = +0.08 (n.s.). (**G**) Donor-level coupling of the anabolic program with DAM module pooled across all donors (single regression, pooled Spearman R). (**H**) Donor-level coupling of the anabolic program with the DAM module; within-Braak regression with the partial Spearman R (controlling for Braak stage).

Because the anabolic program is TREM2-dependent in mice, we asked whether TREM2 expression tracks anabolic capacity in human microglia. Across donors, *TREM2* expression was positively correlated with the anabolic module score at every Braak stage (Fig. 8E). Within DAM-high microglia, TREM2 expression correlated positively with the anabolic score, and this coupling persisted after controlling for Braak stage (Braak I-II: +0.35; Braak III-IV: +0.24; Braak V-VI: +0.34) (Extended Fig. 8E), indicating that the association is not merely a consequence of both TREM2 and anabolic capacity varying with disease stage.

Finally, we examined the relationship between the anabolic and DAM programs across disease progression. The two programs followed opposing trajectories: mean anabolic scores declined while mean DAM scores rose with advancing Braak stage, the trends separating most clearly by Braak V-VI (Fig. 8F). The pooled correlation between the anabolic and DAM scores was near zero (Spearman R = 0.08) (Fig. 8G), reflecting these opposing stage-wise trends. After controlling for Braak stage, however, the anabolic program was positively coupled with both DAM (partial R = 0.23) (Fig. 8H) and *TREM2* (partial R = +0.28, P<1×10⁻⁴) (Extended Fig. 8F) and was uncoupled from the homeostatic module (partial R = +0.08, n.s.) (Extended Fig. 8G). Thus, within a given disease stage, anabolic and DAM activation co-occur, but as pathology advances the anabolic program declines even as the DAM signature is maintained. This mirrors the dissociation observed in the mouse, in which acquisition of the DAM state did not guarantee anabolic competence.

Together, these human data indicate that the microglial anabolic program is conserved across species and that its capacity erodes with AD progression, in association with advancing pathology. The persistence of the DAM signature alongside a declining anabolic program in late-stage human disease demonstrates that DAM signature and biosynthetic competence are separable, and the latter is a more informative correlate of functional microglial resilience.

## Discussion

Microglial activation in Alzheimer’s disease has canonically been defined by the transition to a Disease-Associated Microglia (DAM) state. Because this signature includes phagocytic receptors (e.g., *Clec7a*, *Axl*, *MerTK*), the DAM phenotype has often been treated as synonymous with enhanced phagocytic function. Our findings indicate that this equivalence does not hold. Across single-cell and in situ analyses, the DAM signature and phagocytic competence were dissociable: DAM module scores tracked with pathological burden even in microglia that were metabolically failing, whereas a distinct, TREM2-dependent anabolic program (enhanced protein synthesis coupled to mitochondrial biogenesis) better distinguished functionally engaged cells. We therefore interpret these as molecularly separable programs: the DAM signature may represent a transcriptional stress response to pathology (the ‘demand’ for clearance), whereas the anabolic program reflects the biosynthetic capacity to act on it (the ‘supply’). As amyloid burden rises, demand increases while this supply is progressively suppressed, producing a supply–demand mismatch. This distinction offers one explanation for why transcriptional state alone is often a poor predictor of microglial function. We note that human microglia encompass a broader spectrum of states than the DAM paradigm captures; we position anabolic adaptation as a functional axis within this taxonomy rather than as a replacement for it.

The catabolic burden of Aβ containment was consistently accompanied by an anabolic response. In peripheral macrophages, metabolic states are often dichotomized into glycolysis-driven pro-inflammatory activation versus oxidative-phosphorylation-dependent anti-inflammatory states ^12^. Plaque-associated microglia in our datasets did not fit this binary, instead engaging biomass synthesis and mitochondrial respiration simultaneously, reminiscent of the mTOR-driven expansion of translational and mitochondrial capacity that supports effector function in activated T cells ^18^ ^19^. The functional rationale we propose is proteostatic: continuous internalization of Aβ and cellular debris consumes lysosomal enzymes, membranes, and ATP faster than homeostatic synthesis can replace them, and the observed upregulation of ribosomal and mitochondrial biogenesis may serve to maintain this proteostatic bandwidth. Previous studies have suggested a TREM2-mTOR axis supporting microglial metabolic fitness ^13^. Our data are consistent with this and extend it: the TREM2-dependent anabolic program supports microglial proteome remodeling and mitochondrial renewal, is dissociable from the DAM transcriptional state, and is self-limiting across the amyloid gradient. Within this framework, the synaptic and myelin debris accumulating in *Trem2*^KO^ microglia may reflect not a primary defect in recognition or uptake but a failure of degradative processing, although we cannot exclude contributions from altered uptake or cargo-processing kinetics.

Our metabolic profiling points to impaired mitochondrial renewal as one mechanism that may underlie microglial dysfunction in the absence of TREM2. Our single-cell and in situ data showed a decoupling of organelle mass from function. *Trem2*^KO^ microglia displayed a ‘High Mass / Low Potential’ phenotype, accumulating depolarized mitochondria. We propose that this reflects an impaired ability to renew the mitochondrial network: under chronic phagocytic stress, sustained turnover requires continuous biogenesis, and our in vivo AHA-TOM20 PLA data indicate that TREM2 is required for this biogenic response. We further speculate that, unable to synthesize replacements, *Trem2*^KO^ microglia retain aging, damaged mitochondria (because degrading existing organelles via mitophagy may be disadvantageous if they cannot be replaced), yielding the observed increase in total mass alongside reduced membrane potential, with attendant bioenergetic and oxidative consequences ^20^. We acknowledge that our data only establish the mass/potential decoupling and the TREM2-dependence of biogenesis, but not the causal chain linking them.

Extending these observations toward human disease, we examined microglial anabolic adaptation in an integrated human single-nucleus atlas spanning Braak-staged samples. The anabolic program declined in microglia at advanced Braak stage (V-VI), and remained positively coupled to *TREM2* within disease stage, indicating that it is conserved in human microglia and erodes with disease progression. The mouse X04 gradient and the human Braak gradient thus describe the same load-dependent suppression at two scales, acute phagocytic engagement and cumulative pathology, respectively. Because this analysis is cross-sectional and correlative, and because the change is most robustly stated at the level of the anabolic-high microglial fraction, we interpret it as evidence that reduced anabolic capacity is associated with, rather than demonstrated to cause, advancing pathology. Nonetheless, the cross-species concordance strengthens the relevance of the anabolic axis to human AD.

These observations have potential implications for anti-amyloid immunotherapy. The clinical efficacy of Aβ-targeting antibodies depends in part on the capacity of microglia to perform Fc-receptor-mediated phagocytosis^21^ ^22,23^, which imposes a substantial bioenergetic cost. If the anabolic program that sustains phagocytosis is itself suppressed by phagocytic burden, then sustained pharmacological stimulation could drive microglia toward the biosynthetic exhaustion, in which clearance demand outpaces biosynthetic supply. This raises the hypothesis, to be tested directly, that pairing amyloid immunotherapy with interventions that bolster the anabolic axis (for example, TREM2 agonists or metabolic-cofactor supplementation) could help sustain microglial clearance capacity and extend the therapeutic window. We present this as a hypothesis generated by our findings, not a demonstrated therapeutic principle, and note that it is directly testable by measuring the anabolic signature in immunotherapy-treated tissue.

Several limitations qualify our conclusions. First, our central comparisons rely on TREM2 deletion, which affects multiple microglial functions beyond anabolism; without an orthogonal manipulation that targets the anabolic axis directly, we cannot fully separate a cell-autonomous anabolic requirement from accompanying shifts in population composition, and our mechanistic claims are accordingly associative. Our in situ, per-cell measurements argue against a purely compositional explanation but do not exclude one. Second, the human analysis is cross-sectional and correlative. We have framed the corresponding claims to remain within what these data support.

Together, our findings position TREM2-dependent anabolic adaptation as a spatially resolved, human-relevant correlate of microglial proteostatic competence, and suggest that the biosynthetic capacity to sustain phagocytosis, not transcriptional activation alone, marks functionally resilient microglia, with direct consequences for therapies that rely on that capacity.

## Author Contributions

Conceptualization, [D.L., J.G.]; Methodology, [D.L., J.R.A., W.W., A.M.]; Investigation, [D.L., M.C., S.J., W.W., A.M.]; Formal Analysis, [D.L., W.W., A.M.] (bioinformatics / single-cell and proteomic analysis); Resources, [B.M.S., A.M., J.G.]; Writing – Original Draft, [D.L., J.G.]; Writing – Review & Editing, [D.L., J.G., W.W., A.M., J.R.A., B.M.S.]; Visualization, [D.L.]; Supervision, [J.G.]; Funding Acquisition, [J.G.].

## Acknowledgments

We thank Dr. Takaomi C. Saido in RIKEN Center for Brain Science Institute, Wako, Japan for kindly sharing *App*^NL-GF^ mice. We acknowledge resources from the Campus Microscopy and Imaging Facility (RRID:SCR_025078) and the OSU Comprehensive Cancer Center Microscopy Shared Resource, The Ohio State University. This research was supported by National Institutes of Health Grants R01AG073310 and R21AG075875 (to JG), as well as BrightFocus Foundation Awards for Alzheimer’s research (to JG). The content is solely the responsibility of the authors and does not necessarily represent the official views of the National Institutes of Health.

## STAR METHODS

### KEY RESOURCES TABLE

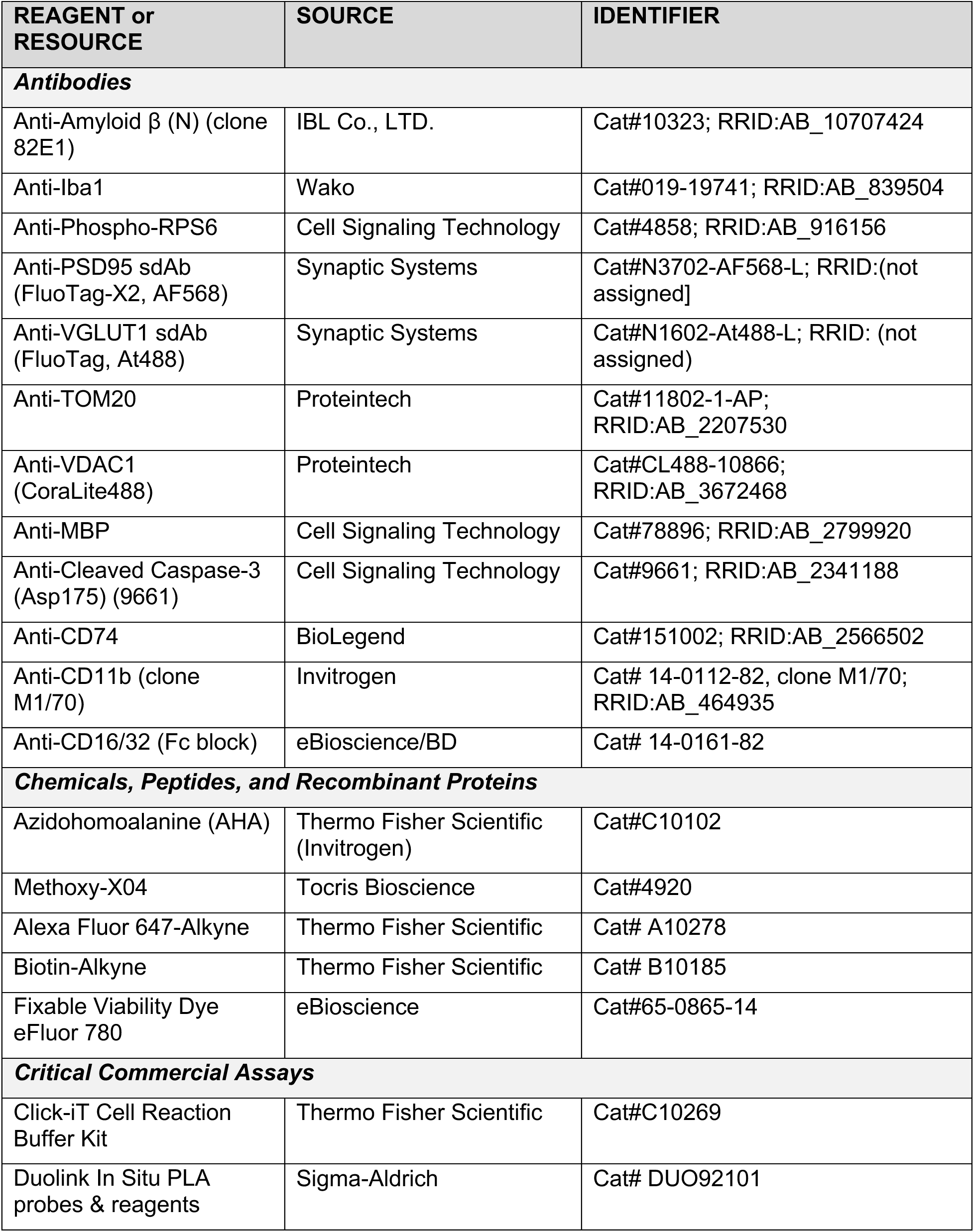
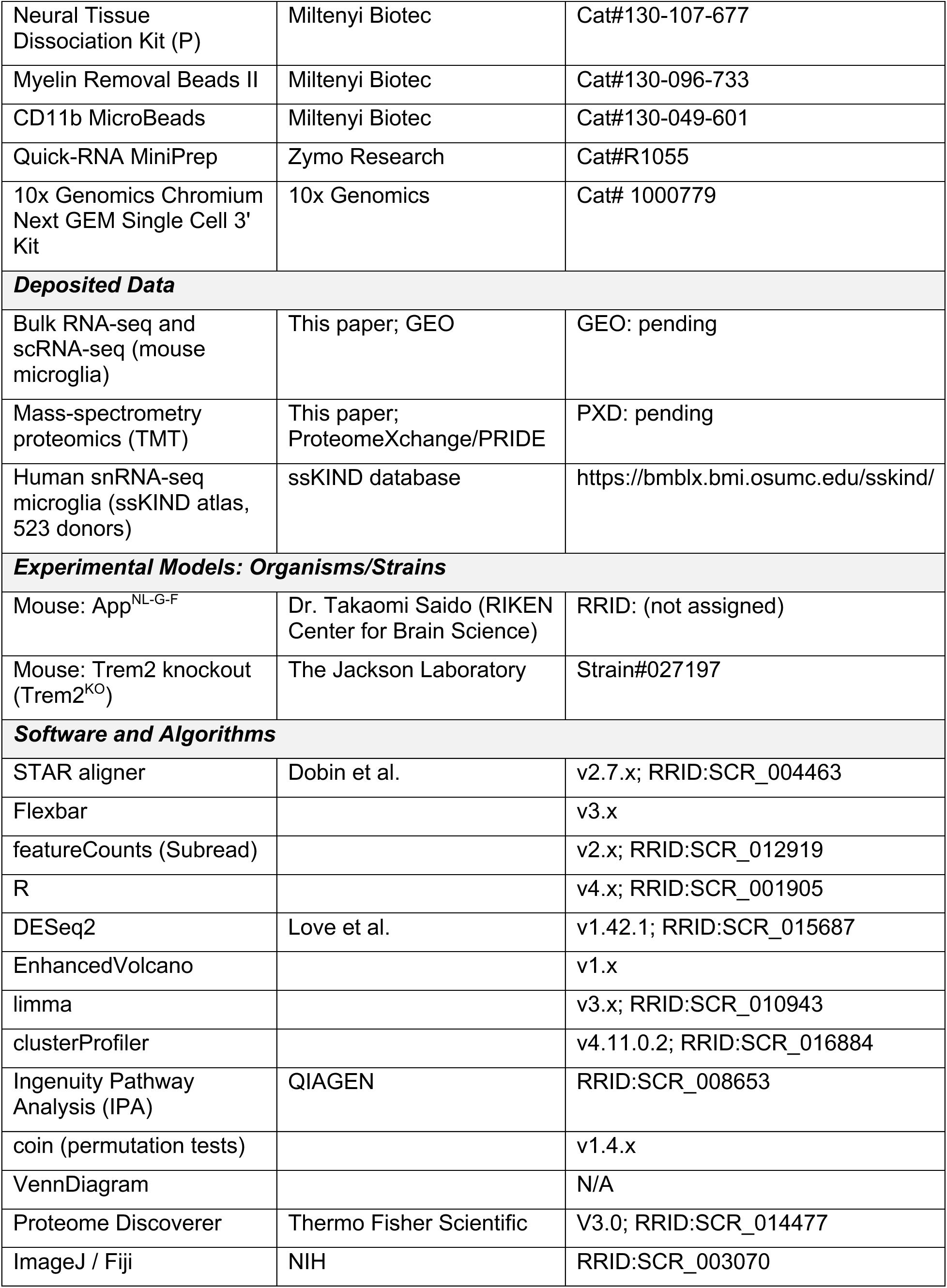
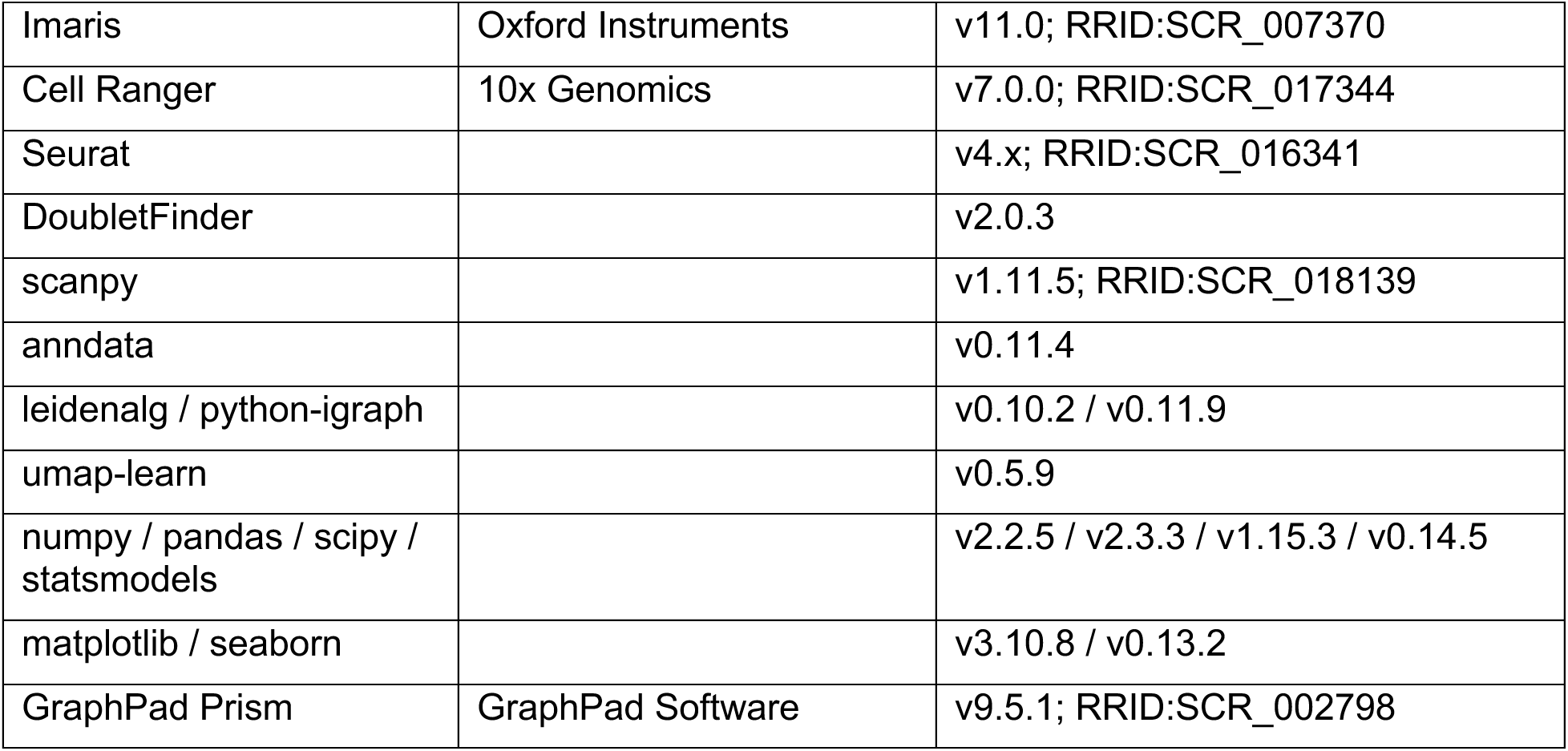

## RESOURCE AVAILABILITY

### Lead Contact

Further information and requests for resources and reagents should be directed to Lead Contact, Jie Gao.

### Materials Availability

This study did not generate new unique reagents. Data and Code Availability

• Bulk RNA-sequencing and single-cell RNA-sequencing data have been deposited at GEO and are publicly available as of the date of publication. The accession number is listed in the Key Resources Table. (GEO pending)

• Mass-spectrometry proteomics data have been deposited to the ProteomeXchange Consortium via the PRIDE partner repository and are publicly available as of the date of publication. The accession number is listed in the Key Resources Table. (PXD pending)

• Human single-nucleus RNA-seq data were obtained from the ssKIND atlas (523 donors) (https://bmblx.bmi.osumc.edu/sskind/)

• This paper does not report original code

• Any additional information required to reanalyze the data reported in this paper is available from the Lead Contact upon request.

## EXPERIMENTAL MODEL AND SUBJECT DETAILS

### Animals

App^NL-G-F^ mice mice were kindly provided by Dr. Takaomi Saido (RIKEN Center for Brain Science) ^24^. Trem2 knockout mice (Trem2^KO^) were obtained from The Jackson Laboratory (Strain #027197). To generate the double-mutant cohort, App^NL-GF^ mice were crossed with Trem2^KO^ mice to produce generate App^NL-G-F^; Trem2^KO^ homozygous for both manipulations. Mice were housed in a pathogen-free barrier facility with a standard 12-hour light/dark cycle and ad libitum access to food and water. Housing density was maintained at maximum 5 mice per cage. Both male and female mice were used for all experiments. All animal procedures were conducted in strict accordance with the National Institutes of Health (NIH) guidelines and approved by The Ohio State University Institutional Animal Care and Use Committee (IACUC).

### Human microglial data analysis

#### Dataset and microglia selection

Human microglia were drawn from the integrated ssKIND single-nucleus atlas ^17^. Cells were retained when annotated as microglia (cell_type = “Microglia”), from donors with a consistent Alzheimer’s disease diagnosis (Disease_consistent = “AD”), and with valid Braak and age metadata. Braak stage was grouped as 0–II, III–IV, V–VI; donor age as 51–78 years and >78 years (the youngest 20–51-year band contained no qualifying AD microglia). The final dataset comprised 204,283 microglia from 523 donors, pooled across brain regions. Raw integer counts were retained for pseudobulk aggregation; the integrated scVI latent space (30 dimensions) was retained for clustering and embedding. Analyses pool donors across age (only Braak stage is distinguished) unless otherwise stated.

## METHOD DETAILS

### In Vivo Metabolic Labeling (FUNCAT)

For labeling of nascent proteins, mice were injected intraperitoneally (i.p.) with 50 mg/kg Azidohomoalanine (AHA; Click Chemistry Tools) dissolved in sterile PBS. Animals were sacrificed 16 hours post-injection. For Flow Cytometry: Dissociated cells were reacted with Alexa Fluor 647-Alkyne (Thermo Fisher Scientific) using the Click-iT Cell Reaction Buffer Kit (Thermo Fisher Scientific, Cat #C10269) according to the manufacturer’s instructions. For Proximity Ligation Assay (PLA): Free-floating brain sections were reacted with Biotin-Alkyne (Thermo Fisher Scientific) using the Click-iT Cell Reaction Buffer Kit prior to antibody incubation.

### Microglial Isolation and Sorting

Adult microglia were isolated using magnetic-activated cell sorting (MACS) as described previously 25. Briefly, mice were anesthetized and perfused transcardially with ice-cold PBS to remove circulating leukocytes. Brains were dissected, chilled on ice, and dissociated using the Neural Tissue Dissociation Kit (P) (Miltenyi Biotec, #130-107-677) and the gentleMACS Tissue Dissociator. Cell suspensions were filtered through a 70 µm cell strainer and centrifuged at 300 × g for 10 minutes. Myelin was depleted using Myelin Removal Beads II (Miltenyi Biotec, #130-096-733) via magnetic separation. The resulting single-cell suspension was used immediately for flow cytometry or further enriched for transcriptomic analysis using CD11b MicroBeads (Miltenyi Biotec, #130-049-601) and LS columns.

### Evaluation of Phagocytic Microglia

To evaluate amyloid load, Methoxy-X04 (X04; Tocris Bioscience, #4920) was prepared by dissolving the compound in DMSO, followed by dilution in a 1:1 mixture of propylene glycol and PBS to obtain a stable yellowish-green emulsion. The solution was prepared freshly and injected i.p. at 10 mg/kg 16 hours prior to tissue harvest. For flow cytometric analysis, dissociated cells were incubated with Fixable Viability Dye eFluor 780 (1:1000, eBioscience) and anti-CD16/32 (Fc Block, 1:200, clone 2.4G2) to exclude dead cells and prevent non-specific binding. Cells were subsequently stained with fluorophore-conjugated antibodies against CD11b (1:500, clone M1/70, Invitrogen), CD45 (1:100, clone 30-F11, Invitrogen), and CX3CR1 (1:50, R&D Systems). Live microglia (CD45int; CD11b+; CX3CR1+) were gated as X04+ or X04-based on fluorescence in the DAPI excitation channel (405 nm). Wild-type animals injected with Methoxy-X04 served as biological negative controls to define gating thresholds.

### In Situ Proximity Ligation Assay (PLA)

In Situ Proximity Ligation Assay (PLA): To visualize nascent mitochondrial proteins in situ, free-floating brain sections from AHA-injected mice were first subjected to the click reaction with Biotin-Alkyne as described above. Sections were extensively washed and incubated overnight at 4 °C with rabbit anti-TOM20 (1:500, 11802-1-AP, Proteintech) and mouse anti-Biotin (1:500, 1D4-C5, BioLegend). PLA probes (Duolink In Situ, Sigma-Aldrich) were applied, and ligation and amplification steps were performed according to the manufacturer’s protocol.

### Bulk RNA-seq and Bioinformatics Analysis

#### Library Preparation and Sequencing

Total RNA from primary microglia was extracted using Quick-RNA miniprep (R1055, Zymo Research). RNA quality was evaluated by TapeStation using high Sensitivity RNA ScreenTape (5067-5579, Agilent). RNA samples with RNA integrity numbers greater than 8 were used for cDNA library construction. RNA seq libraries were prepared using SMART Seq® mRNA LP Kit (Takara Bio) following the manufacturer’s instructions. The qualities of the cDNA library were assessed using TapeStation using High Sensitivity D5000 ScreenTape (5067-5592, Agilent). cDNA library samples were then pooled and sequenced with the HiSeq 4000 System (Illumina) by AZENTA life sciences.

#### Raw Reads Preprocessing and Sequence Alignment

Demultiplexed FASTQ files of bulk RNA sequencing data were aligned to the mouse genome (Mus_musculus.GRCm39) using STAR (version 2.7.10a) ^26^ ^27^. Adapters were trimmed using Flexbar (version 3.5.0.) ^28^. Reads mapped to genomic features were counted using featureCounts (version 2.0.3) ^29^. The count matrix was imported in R (version 4.3.3) for analysis.

#### Quantitative Proteomic Analysis

Quantitative Proteomic Analysis: Brain samples were lysed using 5% SDS and 50mM TEAB buffer, sonicated, cleaned, and quantified via BCA assay. Proteomic analysis was conducted by BGI Genomics Co., Ltd. Briefly, samples were then prepared using the STrap Midi MS sample prep device (Protifi), with each sample containing 2000μg of protein. This preparation involved reduction with dithiothreitol (DTT), alkylation with iodoacetamide (IAM), quenching of the IAM reaction with DTT, and overnight digestion with Trypsin/Lys-C within the STrap device. Peptides were eluted from the STrap, with 60μg per sample dried via SpeedVac and reconstituted in 50% acetonitrile for tandem mass tag (TMT) labeling in 50mM TEAB (pH 8.5). After TMT labeling, samples were pooled, acidified with 1% formic acid, and analyzed for label check on a nano LC-MS/MS system. Following successful label checks, samples were dried, reconstituted in 2% formic acid, desalted using EVOLUTE® EXPRESS ABN (Biotage), and fractionated via offline HPLC into 12 fractions. These fractions were analyzed by LC-MS/MS after being reconstituted with mobile phase A; approximately 5% of each was injected using the TMT method. TMT quantification and identity discovery were performed using Proteome Discoverer 2.5 (Thermo Fisher). False discovery rate (FDR) was calculated based on The Benjamini-Hochberg Procedure. FDR <= 0.05 was considered to be significant.

#### Histological analysis

Histological analysis: Brains were sectioned on a cryostat at 40-mm thickness. For immunofluorescence staining, free-floating sections were blocked with PBS containing 10% normal goat serum (NGS) at room temperature for 30 minutes, incubated with primary antibody in blocking solution (PBS with 1% NGS) at 4°C for 24-48 hours, and then incubated with secondary antibody at room temperature for 2 hours. Sections were mounted on slides with ProLong Diamond (Life Technologies). Images were captured on a ZEISS Axio Observer and/or the Nikon AXR point scanning confocal microscope. 2D Image quantification was performed using ImageJ software. Auto Threshold methods “Otsu” or “Triangle” were used to define the region of interest (ROI). Statistical analyses were conducted using a two-tailed unpaired t-test or one-way ANOVA.

Primary antibodies used in this study are: Human Amyloid β (N) (82E1, IBL Co., LTD.), Iba1 (019-19741, or 011-27991 from Wako Co), Phos-RPS6 (4858S, Cell Signaling), PSD95 sdAb (N3702-AF568-L, Synaptic Systems), VGLUT1 sdAb (N1602-At488-L, Synaptic Systems), TOM20 (11802-1-AP, Proteintech), VDAC1 (CL488-10866, Proteintech), MBP (78896, Cell Signaling), Cleaved Caspase-3 (Asp175) (9661, Cell Signaling), CD74 (151002, Biolegend). All secondary antibodies were purchased from ThermoFisher or Jackson Immunoresearch.

### Image Acquisition and Statistical Analysis

#### 2D Epifluorescence Microscopy and Analysis

Two-dimensional epifluorescence imaging was performed using a ZEISS Axio Observer. This modality was employed for analyses where axial depth was not a primary variable. Image quantification was conducted using ImageJ software (NIH) ^34^, focusing on aggregate metrics per field of view. To ensure unbiased measurement, regions of interest (ROIs) were defined using automated thresholding algorithms (’Otsu’ or’Triangle’, depending on signal-to-noise characteristics). Statistical comparisons between experimental groups were performed using either a two-tailed unpaired t-test or one-way ANOVA, as appropriate, based on the number of groups and data distribution.

### Single-cell Seq and Data Processing

#### Library Construction

Single cell RNA-seq libraries were generated using the 10x Genomics Chromium NEXT GEM Single Cell 3’ Reagent Kit. Briefly, primary microglia isolated from adult mice were loaded onto chromium chips with a capture target of 10,000 cells per sample. Libraries were prepared following the provided protocol and sequenced on an Illumina NovaSeq with a targeted sequencing depth of 50,000-100,000 reads per cell. FASTQ files from sequencing were then used as inputs to the 10X Genomics Cell Ranger pipeline.

#### Read Processing, Quality Control and Filtering

Gene expression matrices were generated with the Cell Ranger Pipeline (v7.0.0; 10x Genomics) and aligned to the Mouse (mm10) reference transcriptome. The resulting digital gene expression matrix was filtered, normalized, and clustered using R version 4.2.0 and Seurat version 4.1.1 ^39^. Genes that are expressed in less than 10 cells, and cells with greater than 5% of reads mapped to mitochondrial genes, or with less than 1500 features and 3000 UMIs were removed.

## QUANTIFICATION AND STATISTICAL ANALYSIS

### Differential Expression and Principal Component Analysis (PCA)

Differential gene expression analysis was performed using the DESeq2 package (version 1.42.1) ^30^ in R. Raw count data were filtered to remove genes with low library representation (total counts < 10). For basic comparisons, a single-factor generalized linear model was used. In experiments involving Batch/Sex/Time, a multi-factor model was implemented to control for confounding variables. DESeq2’s internal median-of-ratios method was used for normalization. To control the false discovery rate (FDR), p-values were adjusted using the Benjamini-Hochberg procedure. Genes were considered significantly differentially expressed if they reached an adjusted p-value < 0.05. Differential expression results were visualized using volcano plots generated by the EnhancedVolcano package (version 1.20.0) ^31^, with significance defined as an adjusted p-value < 0.05.

PCA was performed on the transformed counts extracted from DESeq2. If a multi-factor design was used for DEA to measure the effect of the genotypes controlling for batch differences, the PCA was plotted with batch variation removed by using the removeBatchEffect() function from limma (version 3.54.2) ^32^.

### Functional Enrichment and Pathway Analysis

#### IPA Core Analysis

Core Canonical Pathway Analysis was performed by QIAGEN’s Ingenuity® (IPA®, QIAGEN Redwood City, www.qiagen.com/ingenuity). Complete lists of DEGs and DAPs, along with their log2 fold change expression values and FDR were inputted into IPA for identifying canonical pathways, biological functions, and upstream regulators using a cutoff of FDR < 0.05. The p-value of overlap, calculated using the right-tailed Fischer’s Exact Test with a statistical threshold of 0.05, is used to indicate the probability of association of molecules from test dataset with the canonical pathway by random chance alone. A positive or negative regulation z-score value indicates that a function is predicted to be activated or inhibited. No activity prediction by IPA results in ineligible z-score which is represented by grey bars.

#### IPA Comparison Analysis

To identify biological pathways modulated across the different XO4 subgroups of microglia within specific genotype, differentially expressed genes (DEGs) from the Low, Medium, and High groups from each genotype--each compared against a common baseline control of XO4-dataset--were first processed through individual Core Analyses in IPA. Subsequently, the results of contrast within each genotype were integrated using the IPA Comparison

Analysis platform to juxtapose the canonical pathway profiles across all groups of the specific genotype. Findings were visualized as a heatmap where color intensity represents the activation z-score (grey indicates an unpredictable directional trend). To denote statistical confidence, pathways failing to reach the significance threshold (e.g., Fisher’s exact test p-value > 0.05) were marked with an asterisk (*). Pathways were further grouped and annotated by their broader functional categories to identify overarching biological themes.

#### Gene Set Enrichment Analysis (GSEA)

Using the R package clusterProfiler ^33^ (version 4.11.0.2, genes or proteins were ranked by values of log2 Fold Change and −log10(p-value) ∗ sign(log2FoldChange) respectively to form the ranking metrics for transcriptomic and proteomic datasets. Enrichment was conducted against the Gene Ontology (GO) database, specifically targeting Biological Process (BP) and Cellular Component (CC) categories. Results were considered statistically significant at an adjusted p-value < 0.05. Results were visualized using dot plots, where selected GO terms were segregated by their direction of regulation (activated vs. suppressed) based on the Normalized Enrichment Score (NES). The color scale represents the Benjamini-Hochberg adjusted p-value (q-value) to indicate statistical significance, while the size of each dot corresponds to the gene count (the number of genes from the dataset coregulated within a specific GO term).

#### Gene-concept Network

To reveal the molecule-level information associated with the significant pathways of interest, we constructed gene-concept networks using the cnetplot() function within the clusterProfiler package in R. 3D Confocal Imaging and Hierarchical Statistical Modeling

High-resolution three-dimensional image stacks were acquired using the Nikon AXR point scanning confocal microscope to quantify the morphological and biochemical properties of Iba1-positive (Iba1+) regions of interest (ROIs). Due to the high density and clustering of microglia in the App^NL-G-F^ mice, individual ROIs were defined as distinct morphological units rather than individual cells.

All 3D reconstructions and quantitative analyses were performed using Imaris (v11.0; Oxford Instruments, Zurich, Switzerland), a multidimensional image analysis software. Iba1+ surfaces were generated using a machine learning-based thresholding algorithm to ensure objective and consistent segmentation of complex microglial morphologies. To quantify the volumetric colocalization of target markers within Iba1+ ROIs, marker surfaces were generated via manually defined thresholds, allowing for the correction of staining-specific background noise. Volumes were converted to voxel counts prior to calculating the overlapping voxel ratio and performing model fitting. To assess protein expression levels, the mean fluorescence intensity of the specific target marker channel was extracted directly from the reconstructed Iba1+ surface.

To assess the impact of AD pathology, Iba1+ ROIs were spatially categorized based on their interaction with amyloid pathology. An ROI was classified as Plaque-Associated Microglia (PAM) if any portion of the Iba1+ surface, including processes or the soma, was in direct contact with a plaque cluster. All other ROIs were designated as Non-Plaque-Associated Microglia (NPAM).

To account for the hierarchical structure of the 3D data (multiple ROIs nested within individual imaging fields), we employed a Generalized Linear Mixed Model (GLMM) framework, allowing us to treat the animal or image stack as a random effect, thereby controlling for intra-subject variation and ensuring a more accurate estimation of the fixed effects of the genotypes, Plaque Status (PAM vs. NPAM), and their interaction term.

All statistical analyses were conducted in R using the glmmTMB ^35^ and emmeans ^36^ packages. The choice of statistical model was tailored to the distribution and mathematical constraints of each quantified metric. For the volumetric occupancy of target markers within Iba1+ ROIs, a beta-binomial GLMM (logit link) was selected to account for the bounded nature of proportional data and to adjust for overdispersion derived from biological variability. Continuous measurements, such as mean intensities, were modeled using a Gamma distribution (log link).

Model fit was rigorously assessed using the DHARMa package ^37^, utilizing a simulation-based approach to verify distribution assumptions, dispersion, and residual patterns. Estimated Marginal Means (EMMs) and 95% confidence intervals (CIs) were back-transformed from the link scale to the response scale for reporting. Pairwise comparisons were performed as planned contrasts between genotypes and plaque conditions.

#### Microglial Cargo-Inclusion Quantification and Non-Parametric Analysis

Following 3D confocal acquisition as described above, we performed a targeted quantification of microglial cargo inclusions. Inclusion density was quantified via a dual-blind system: Aβ plaque areas were manually delineated by an analyst blinded to the inclusion-marker channels (PSD95 and MBP), and inclusions were subsequently counted within these regions by a second independent analyst.

Inclusion densities were calculated as the number of inclusions per 105 μm3 of plaque volume to ensure human-readable scaling. Due to the distinct biological nature of the groups—where the control (AK) genotype exhibited “structural zeros” (near-total absence of symptoms) and the treatment (AKTK) genotype showed a robust, high-variance phenotype—standard count-based Generalized Linear Mixed Models (GLMMs) failed to reach mathematical convergence. To address the complete separation and non-normal distribution of the data, a non-parametric Wilcoxon rank-sum test with exact permutation and rank transformation using the R package coin (version 1.4.3) ^38^ was employed to account for the high frequency of tied zeros in the control group. Inclusion density distributions were visualized in violin plots, with jittered points indicating individual plaque ROIs. Medians and interquartile ranges (IQRs) are indicated by dots and bars, respectively. Statistical significance is reported as p values based on the exact permutation test.

#### Initial Normalization and Doublet Removal

We performed an initial normalization of post-QC dataset in Seurat to stabilize variance and did not regress out variation associated with percent.mito or percent.rb due to the metabolic relevance mitochondrial and ribosomal gene expression features, despite observed differences between these fractions. DoubletFinder version 2.0.3 ^40^ was used to identify false-negative Demuxlet classifications caused by doublets formed from cells with identical SNP profiles, and an average of 10% of cells per sample were confidently predicted as doublets and removed. In total, 26,096 cells were identified as putative singlets and retained for downstream analysis.

#### SCTransform Normalization, Integration, and Clustering

The gene expression matrix was normalized and scaled using the Seurat function SCTransform which also identifies the most variable genes, of which the top 3,000 were used for dimensionality reduction. Four samples were integrated to correct for any potential library batch effect by using the Seurat functions FindIntegrationAnchors and IntegrateData based on reciprocal PCA with n = 5 neighbors (k.anchor). Integrated matrix was used for downstream analysis. Cells were clustered using the Louvain algorithm based on the first 20 principal components with a resolution of 0.3. The Uniform Manifold Approximation and Projection (UMAP) ^41^ was used for non-linear reduction and two-dimensional data visualization.

#### Cluster Annotation, Subsetting, and Reclustering

Cell-type annotations were assigned to each cluster based on two levels of evidence. First, the Seurat function FindAllMarkers was used to identify cluster marker genes based on one-versus-all Wilcoxon rank sum differential expression tests for each cluster. Second, cell-type identities were predicted by comparing transcriptomic profiles to a curated panel of marker genes derived from a previously published single-cell RNA sequencing dataset of the App^NL-G-F^ mouse brain ^42^. Based on this classification, we retained 25,776 cells identified as putative microglia for downstream analyses, while non-microglial cell types were excluded. For reclustering putative microglia, we applied SCTransform normalization, recomputed PCA and used the top 20 PCs for dimensionality reduction by UMAP, followed by unbiased clustering using the Seurat function FindNeighbors with the resolution of granularity set to 0.2. This led to the identification of 7 clusters each representing a microglial state defined by unique or transitory profiles.

#### Differential Gene Expression Analysis

Differentially expressed genes of specific cell states were found by applying the Seurat function FindAllMarkers for overall DE and FindMarkers for side-by-side comparisons. Genes with adjusted p values (using a Bonferroni correction) < 0.05 were considered significantly differentially expressed. Canonical Pathways Analysis by IPA was used to test for gene sets enriched in DE genes.

#### Gene Module Scoring and Signature Analysis

To compare the transcriptomic profile of our clusters with previously described microglial states, we calculated module scores using the AddModuleScore function in Seurat. Signatures were defined based on published marker lists for homeostatic microglia (HM) markers (Tmem119, P2ry12, Cx3cr1), activated response microglia (ARM) markers (Apoe, Cst7, Itgax, Lpl, Spp1, Gpnmb, Dkk2, Cd74, H2-Aa, H2-Ab1), transiting response microglia (TRM) markers (Apoe, Cst7, Itgax, Cd74, H2-Aa, H2-Ab1), interferon response microglia (IRM) markers (Ifit2, Ifit3, Ifitm3, Oasl2, and Irf7), cycling and proliferating microglia (CPM) markers (Top2a, Mcm2, Tubb5, Mki67, Cdk1), a Ribosomal Microglia signature (Tpt1, Rps3a, Rpl13, Rps23), and Disease-associated Mciroglia (DAM) markers (Cd9, Apoe, Trem2, Tyrobp, Cd63, Lgals3, Axl, Spp1, Cstb, Ctsd, Lpl, Itgax, B2m, Cst7, Gpnmb, Igf1, Irf8, Fth1, Lyz2, Ccl3, Ccl6, Timp2) in App^NL-G-F^ mice as curated in previous publication^43^.

#### Assessment of Signature-level Enrichment across Cell States

To assess the distribution of gene expression across the identified transcriptomic states in our dataset (resolution 0.3), we generated stacked violin plots using the Seurat package. This visualization displays the log-normalized expression levels of representative marker genes, which were grouped along the y-axis into modules corresponding to curated microglial signatures and canonical pathways identified via Ingenuity Pathway Analysis (IPA). Individual cell clusters, denoted by their index numbers, are arrayed along the x-axis.

#### Multi-omics Integration and Comparative Analysis

To elucidate the regulatory landscape across different molecular layers, we performed a comparative and integrative analysis of the transcriptomic (RNA-seq) and quantitative proteomic profiles from primary microglia isolated from 9-month-old and 3-month-old App^NL-G-F^ and APP^NL-G-F^;Trem2^KO^ mice.

#### Direct Profile Comparison

We first assessed the direct correspondence between identified transcripts and proteins. Initial comparison was conducted by mapping identified transcripts to their corresponding proteins. A Venn analysis was utilized to determine the overlap between the two datasets, identifying molecules consistently regulated at both levels as well as those uniquely detected within a single “omic” layer. The resulting intersections were visualized as a Venn diagram using the VennDiagram package (version 1.7.3) ^44^ in R. Integrative Pathway Analysis

To identify biological processes consistently active across both molecular layers, we employed ActivePathways, an integrative method that uses directionality and significance estimates of molecules to identify significantly enriched pathways by combining evidence from multiple omic sources ^45^.

Integration metrics. P-values and log2 fold-change (log2FC) values derived from the 9-month-old versus 2-month-old comparisons were processed using the ActivePathways framework.

Statistical evidence from the transcriptomic and proteomic datasets was integrated via Data-driven P-value Merging (DPM), with Brown’s method utilized as a robust reference for determining combined significance. To identify biologically convergent profiles, a weighted constraint vector [mRNA=1,protein=1] was applied to prioritize genes exhibiting direct, concordant associations between the two molecular layers. Conversely, genes with conflicting directional signals or those failing to meet the integration criteria were penalized, ensuring the final pathway enrichment was driven by consistent cross-omic evidence. The relationship between the two molecular layers was visualized using a concordance scatter plot.

Integrative pathway enrichment analysis. Functional enrichment was performed using the ActivePathways R package to identify biological processes significantly represented across the integrated datasets. The analysis utilized a gene list ranked by merged P-values derived from directional data integration. To determine optimal enrichment of Gene Ontology (GO) terms, a ranked hypergeometric test was applied. The gene set collection (m5.go.v2023.2.Mm.symbols.gmt) was filtered to include only pathways containing between 10 and 500 annotated genes in order to minimize biases from excessively specific or overly generic terms. Significant GO terms were defined using a Holm Family-Wise Error Rate (FWER) < 0.05. To facilitate biological interpretation, unique and significant GO terms were visualized via bar charts with the x-axis representing the −log10(adjusted P-values) and the y-axis listing the specific GO terms which were further grouped by their broader functional categories.

## Statistical Analysis

Statistical analyses were performed using GraphPad Prism 9 software (v9.5.1; GraphPad Software, San Diego, CA, USA). Data are presented as mean ± standard error of the mean (SEM). Pairwise comparisons were analyzed using two-tailed unpaired t-tests. Multiple comparisons were analyzed using one-way or two-way analysis of variance (ANOVA) followed by Tukey’s or Sidak’s post-hoc tests. A P value < 0.05 was considered statistically significant.

### Gene modules

Four gene modules were scored. The anabolic module combines oxidative-phosphorylation subunits (ATP5E, COX4I1, NDUFA1, NDUFB9, NDUFS8, UQCRB, COX7B) and protein-translation genes (RPS3A, RPS9, RPL10, RPL13, RPL18A, EIF3F, EEF1A1, MRPL42, MRPS14). The DAM (disease-associated microglia) module comprised SPP1, APOE, CD9, ITGAX, GPNMB, CD68, CST7, LPL, LGALS3; the homeostatic module comprised P2RY12, CX3CR1, TMEM119, HEXB. A housekeeping track (ACTB, GAPDH, B2M) was scored only as an RNA-quality control.

### Donor pseudobulk and module scores

For each donor, raw counts of its microglia were summed gene-wise, normalized to counts per million (CPM) and log2(CPM+1) to give a donor pseudobulk profile. A donor module score is the mean, across the genes of a module, of the per-gene z-score of log2CPM, with the z-score computed across donors. This donor-level design (one value per donor) avoids the inflated significance produced by treating individual cells as independent replicates (pseudoreplication).

### Cell-level scores, re-clustering and embedding

For the single-cell visualizations (UMAP), per-cell module scores were computed as the mean across module genes of the per-gene z-score of log-normalized expression, with z computed across cells. Microglia were re-clustered on the integrated scVI latent restricted to these cells: a k-nearest-neighbour graph (k = 15) was built on the 30-dimensional latent, Leiden clustering at resolution 0.4 yielded 11 subclusters, and a UMAP was re-embedded from the same graph. A module-gene dot plot summarizes mean expression (z across clusters) and the fraction of expressing cells per subcluster.

### TREM2 and DAM stratification

A cell was scored TREM2-positive if it carried at least one TREM2 count and TREM2-negative otherwise; per donor, the TREM2-positive fraction and separate TREM2-positive / TREM2-negative pseudobulk profiles were computed. DAM-high cells were defined as the top tercile of the cell-level DAM score (computed across all cells); a donor pseudobulk restricted to DAM-high cells was used for the within-DAM analyses. TREM2 is a low-abundance transcript in droplet single-nucleus data, so the TREM2-positive fraction reflects detection sensitivity and is interpreted comparatively (across Braak stage) rather than as an absolute positivity rate.

### Statistics

Anabolic decline across Braak was tested by ordinary least squares on donor pseudobulk (score ∼ age + Braak + age×Braak); median genes per cell, percent mitochondrial reads and the housekeeping module were modelled identically as RNA-quality controls. Pairwise comparisons between adjacent Braak stages used the Mann–Whitney U test (donor-level, unpaired).

Associations between module scores and TREM2 expression were quantified by Spearman’s rank correlation (ρ, reported on figures as R) overall, within each Braak stage, and within DAM-high cells; rank correlation was used because donor-level scores contain outliers to which Pearson correlation is sensitive. To distinguish coupling that is independent of disease stage from the across-stage trend (a Simpson’s paradox in which the pooled anabolic–DAM correlation is masked by the opposing Braak gradients of the two programs), partial Spearman correlations controlling for Braak stage were computed. The DAM/anabolic dissociation was summarized by per-stage mean module scores; the change in the within-stage anabolic–DAM correlation between early (Braak 0–II/III–IV) and late (Braak V–VI) disease was tested with a Fisher z comparison. To ask whether the donor-level anabolic decline is compositional or per-cell, cells were labelled anabolic-high at a single global threshold (top 30% of the cell-level anabolic score across all cells); per donor we computed the fraction of anabolic-high cells and the mean score within them, and the Braak 3–4 to 5–6 change in the donor mean was split into a compositional and a per-cell component by shift-share decomposition. All tests were two-sided.

### Software

All analyses were performed in Python 3.10.6. Single-nucleus data handling, neighbor-graph construction and UMAP embedding were carried out with scanpy (v1.11.5) ^46^ and anndata (v0.11.4) ^47^; subclusters were identified by Leiden clustering using leidenalg (v0.10.2) ^48^ with python-igraph (v0.11.9), and embeddings were computed with umap-learn (v0.5.9) ^49^. Statistical analyses of microglia used numpy (v2.2.5), pandas (v2.3.3), scipy (v1.15.3) ^50^ and statsmodels (v0.14.5). Figures were generated with matplotlib (v3.10.8) ^51^ and seaborn (v0.13.2) ^52^.

**Table 1.** Cohort characteristics of the human AD microglia dataset, by Braak stage. Data comprise 523 donors (313 female / 210 male) drawn from 14 single-nucleus RNA-seq studies.

| **Characteristic** | **Braak 0–II** | **Braak III–IV** | **Braak V–VI** | **Total** |
| --- | --- | --- | --- | --- |
| **Donors, n** | 34 | 223 | 266 | 523 |
| **Age, years- mean (range)** | 81.2 (64–96) | 89.2 (68–108) | 83.7 (56–101) | 85.9 (56–108) |
| **Sex, F / M** | 21 / 13 | 130 / 93 | 162 / 104 | 313 / 210 |
| **Platform (snRNA / scRNA)** | 34 / 0 | 223 / 0 | 266 / 0 | 523 / 0 |
| **Microglia, n (nuclei)** | 8,445 | 53,609 | 142,229 | 204,283 |

*All donors are single-nucleus RNA-seq. Braak stage groups: 0–II, III–IV, V–VI.*

## Extended Figure Legend

**Extended Figure 1.**
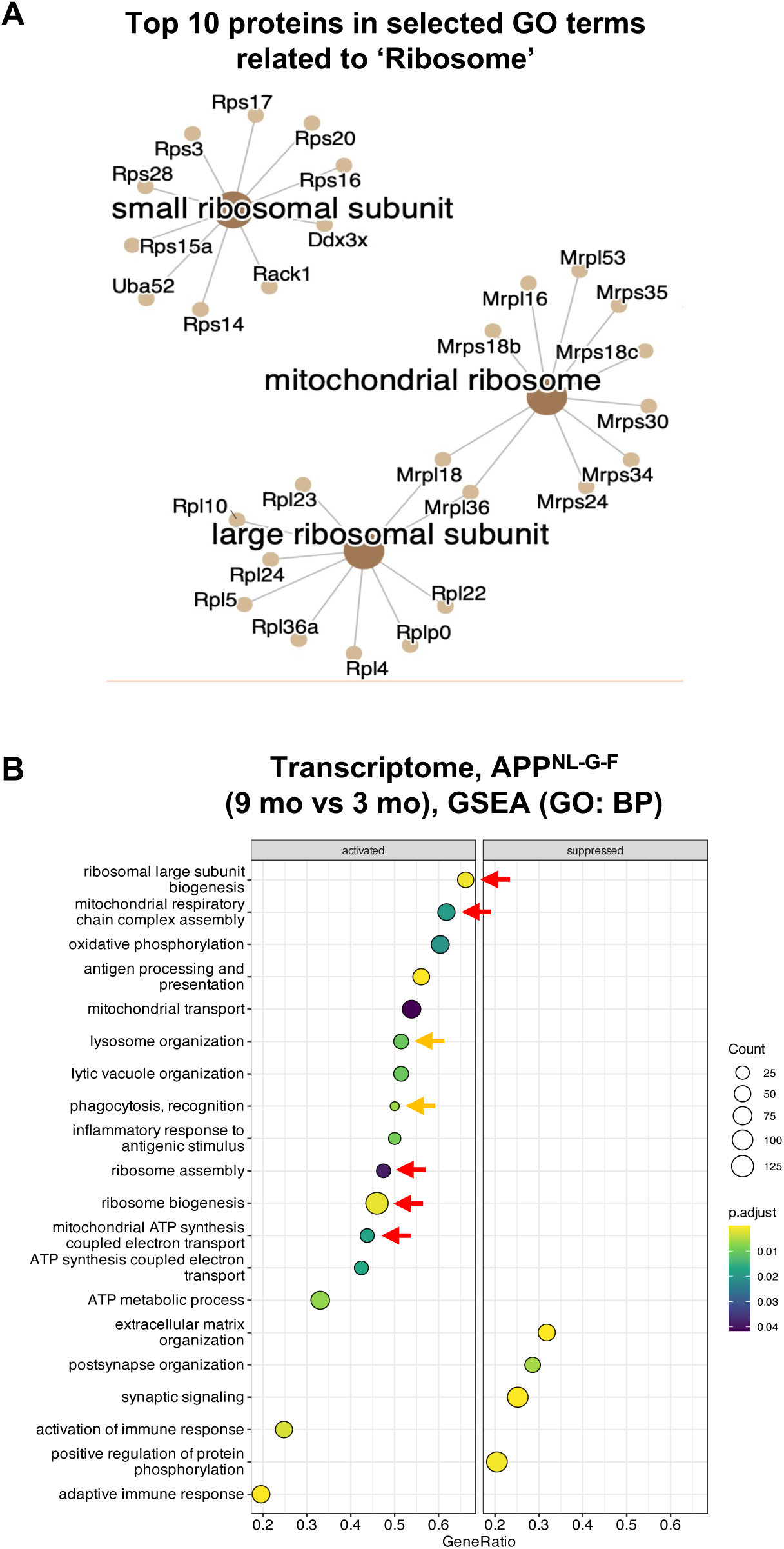
**(A)** Gene-concept network plots display the top 10 differentially abundant proteins associated with the Gene Ontology (GO) terms ‘small ribosomal subunit’, ‘large ribosomal subunit’, and ‘mitochondrial ribosome’. These proteins are significantly enriched in 9-month-old *App^NL-G-F^* microglia compared to 3-month-old controls. Visualization was generated using the Kamada–Kawai layout algorithm. **(B)** Gene Set Enrichment Analysis (GSEA) dot plots of Biological Process (BP) terms representing the transcriptome of *App^NL-G-F^* microglia. Red arrows indicate the enrichment of anabolic signatures and orange arrows indicate catabolic signatures enriched in 9-month-old *App^NL-G-F^* microglia compared to 3-month-old counterparts.

**Extended Figure 2.**
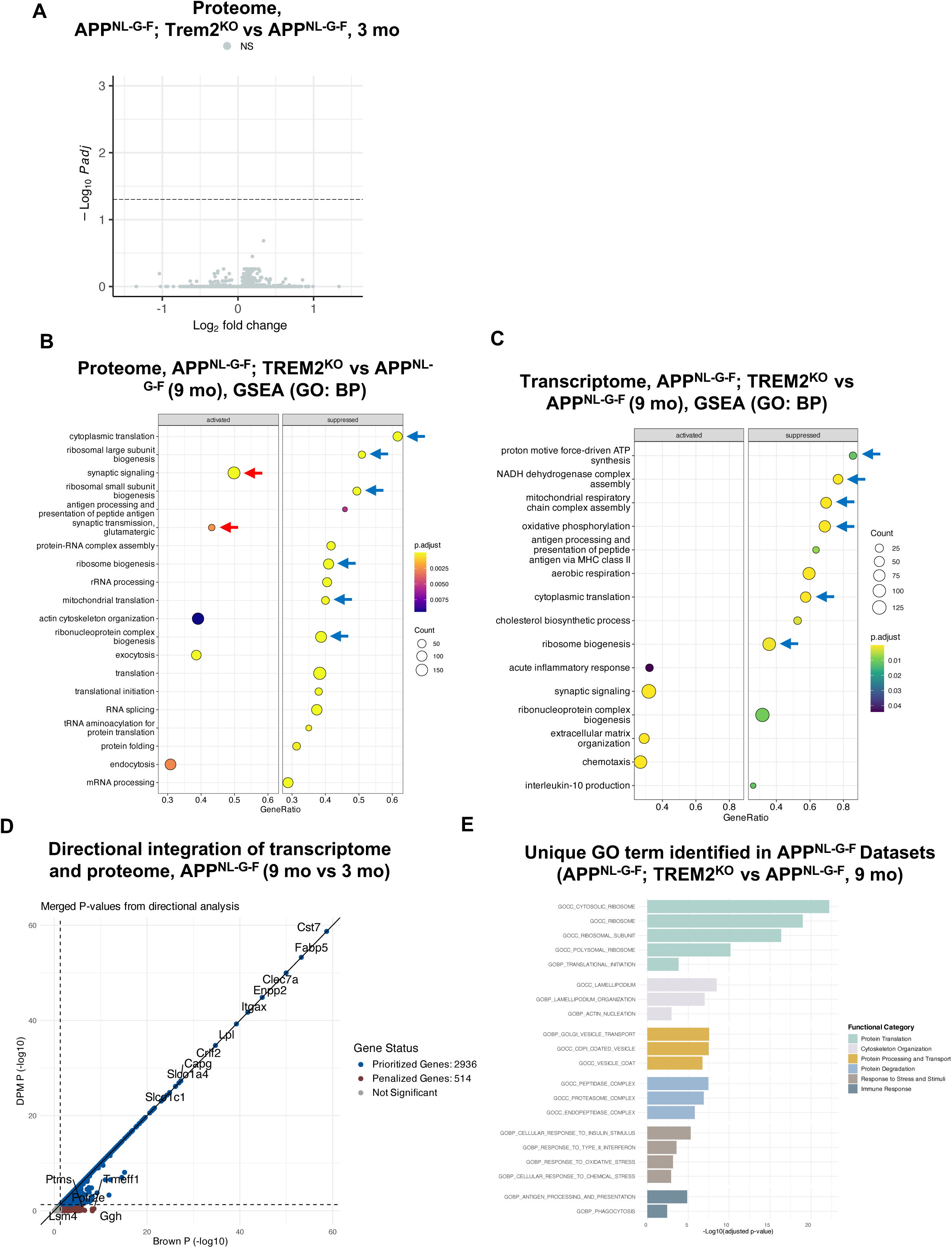
**(A)** Volcano plot showing the absence of differentially abundant proteins between *App^NL-G-F^;Trem2^KO^* and *App^NL-G-F^* microglia at 3 months of age (adjusted *P* < 0.05). **(B, C)** GSEA of BP terms representing the proteome and transcriptome of *App^NL-G-F^;Trem2^KO^*microglia. Dot plots illustrate enriched pathways in the proteome **(B)** and transcriptome **(C)** of *App^NL-G-F^;Trem2^KO^*microglia compared with *App^NL-G-F^* counterparts at 9 months of age. Blue arrows indicate suppressed’ribosomal’ signatures. **(D)** Correlation analysis of transcriptome and proteome directional integration using ActivePathways. Scatter plot illustrating merged *P*-values calculated via directional analysis (DPM, y-axis) and non-directional analysis (Brown’s method, x-axis) in *App^NL-G-F^* microglia (9 months vs 3 months). Prioritized genes with directionally consistent changes across datasets are positioned on or near the diagonal (blue), whereas genes with conflicting directional changes are penalised and located further below the diagonal (brown). **(E)** Multi-omics pathway enrichment of *App^NL-G-F^*microglia. Bar chart illustrating unique GO terms identified in *App^NL-G-F^* microglia but absent in *App^NL-G-F^;Trem2^KO^* microglia datasets. Multi-omics data fusion was performed using ActivePathways, with merged significance estimates generated via the directional-value merging (DPM) method. These prioritized terms are associated with proteostasis and are grouped into six functional categories (indicated by color). Significance was determined using merged *P*-values as input for enrichment analysis.

**Extended Figure 3.**
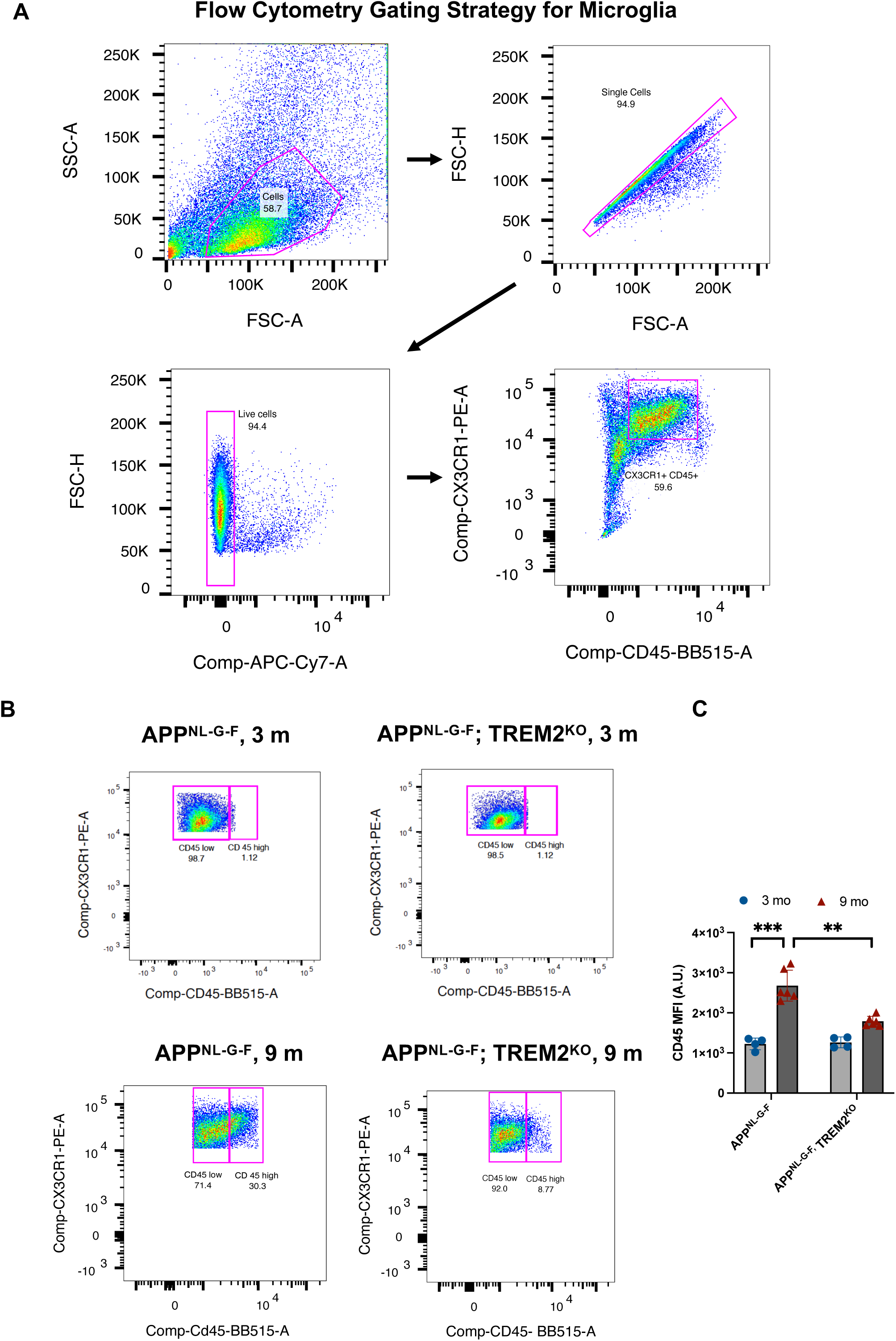
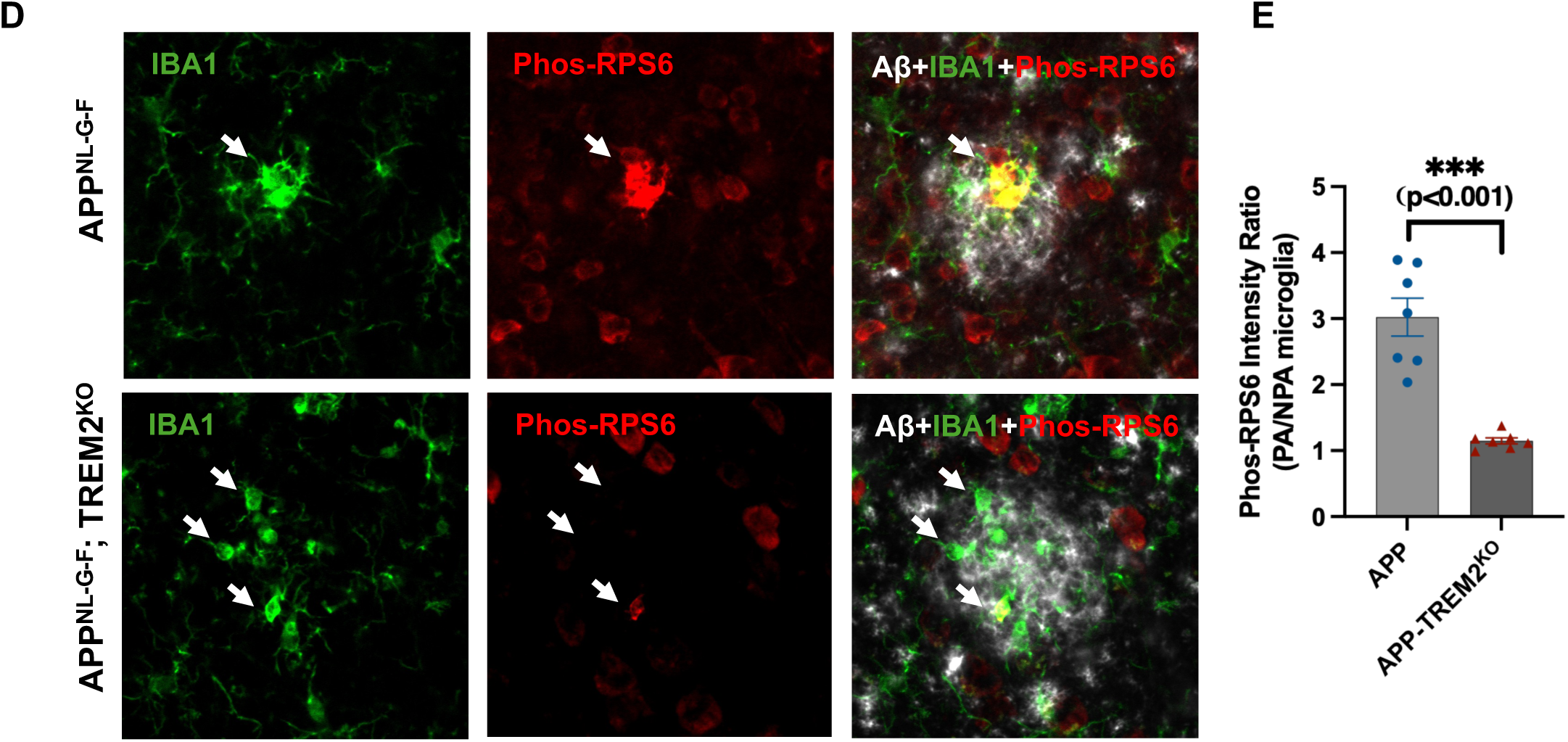
**(A)** Representative flow cytometry gating strategy used to isolate live, singlet microglia (CX3CR1^+^ CD45^+^) from dissociated brain tissue. **(B)** Representative flow cytometry plots illustrating the temporal expansion of the CD45^high^ microglial population from 3 to 9 months of age in *App^NL-G-F^* mice. Note that this expansion is severely blunted in *App^NL-G-F^;Trem2^KO^* mice. **(C)** Quantification of CD45 Mean Fluorescence Intensity (MFI) in primary microglia isolated from *App^NL-G-F^* mice and *App^NL-G-F^;Trem2^KO^* mice at 3 and 9 months of age. **(D)** Representative confocal images of phosphorylated ribosomal protein S6 (p-RPS6, Ser235/236) in plaque-associated microglia from 9-month-old *App^NL-G-F^* and *App^NL-G-F^;Trem2^KO^*mice, co-stained for Iba1 and Aβ. **(E)** Quantification of p-RPS6 intensity per microglial cell. n = 6-8 mice per group. \*\*\**P* < 0.001 by two-tailed unpaired t-test.

**Extended Figure 4.**
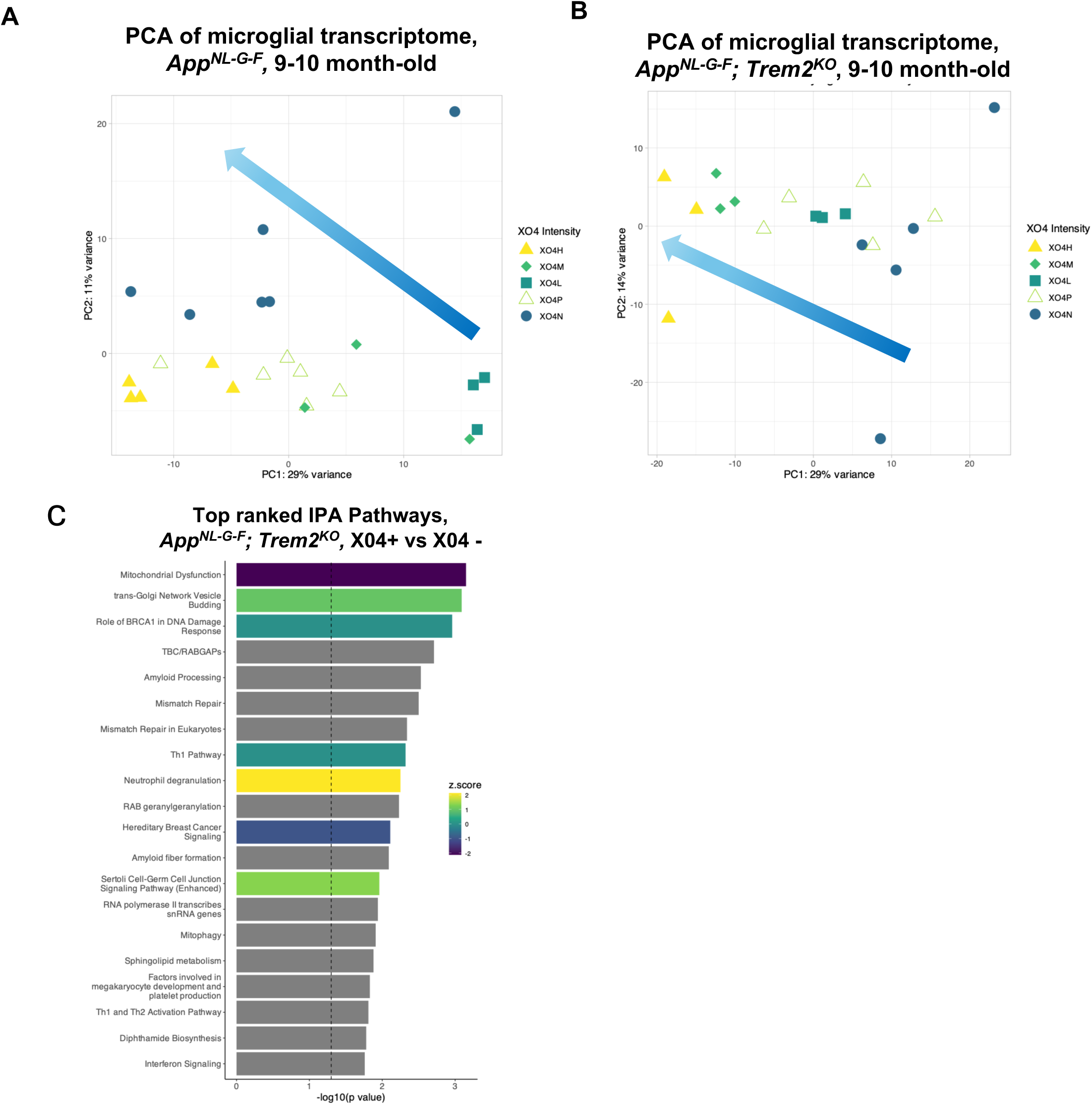
**(A,B)** Distinct transcriptional states of microglia associated with X04 intensity which represents varying degrees of Aβ load in 9-month-old *App^NL-G-F^* and *App^NL-G-F^;Trem2^KO^* microglia. **(A)** PCA of bulk RNA-seq of microglia isolated from 9-month-old *App^NL-G-F^* mice (X04H, n = 5; X04M, n = 3; X04L, n = 3; X04P, n = 6; X04N, n = 6). **(B)** PCA of bulk RNAseq of microglia isolated from 9-month-old *App^NL-G-F^;Trem2^KO^* mice (X04H, n = 3; X04M, n = 3; X04L, n = 3; X04P, n = 5; X04N, n = 5). The arrow (blue to light blue gradient) denotes the spectrum of X04 intensity, ranging from low (right) to high (left). **(C)** IPA Canonical Pathway Analysis comparing X04^+^ versus X04^-^ microglia from 9-month *App^NL-G-F^;Trem2^KO^* mice. Bar chart depicts the top 20 significantly enriched canonical pathways in X04^+^ microglia (*Fisher’s exact test p-value* < 0.05). Note that anabolic adaptation was completely abolished in TREM2-deficient microglia. Pathways are ranked by their enrichment significance, represented as −log10(*P*-value).

**Extended Figure 5.**
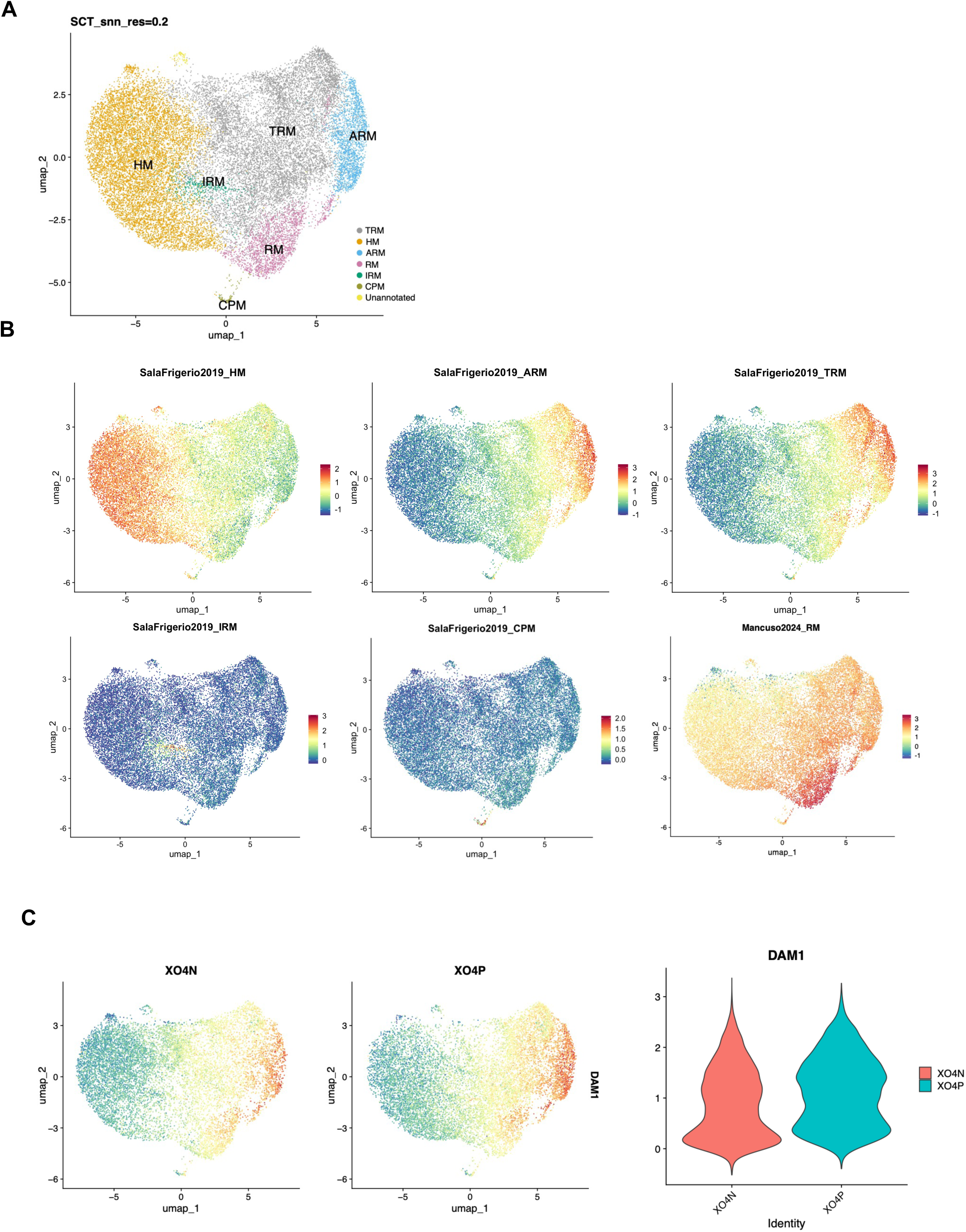
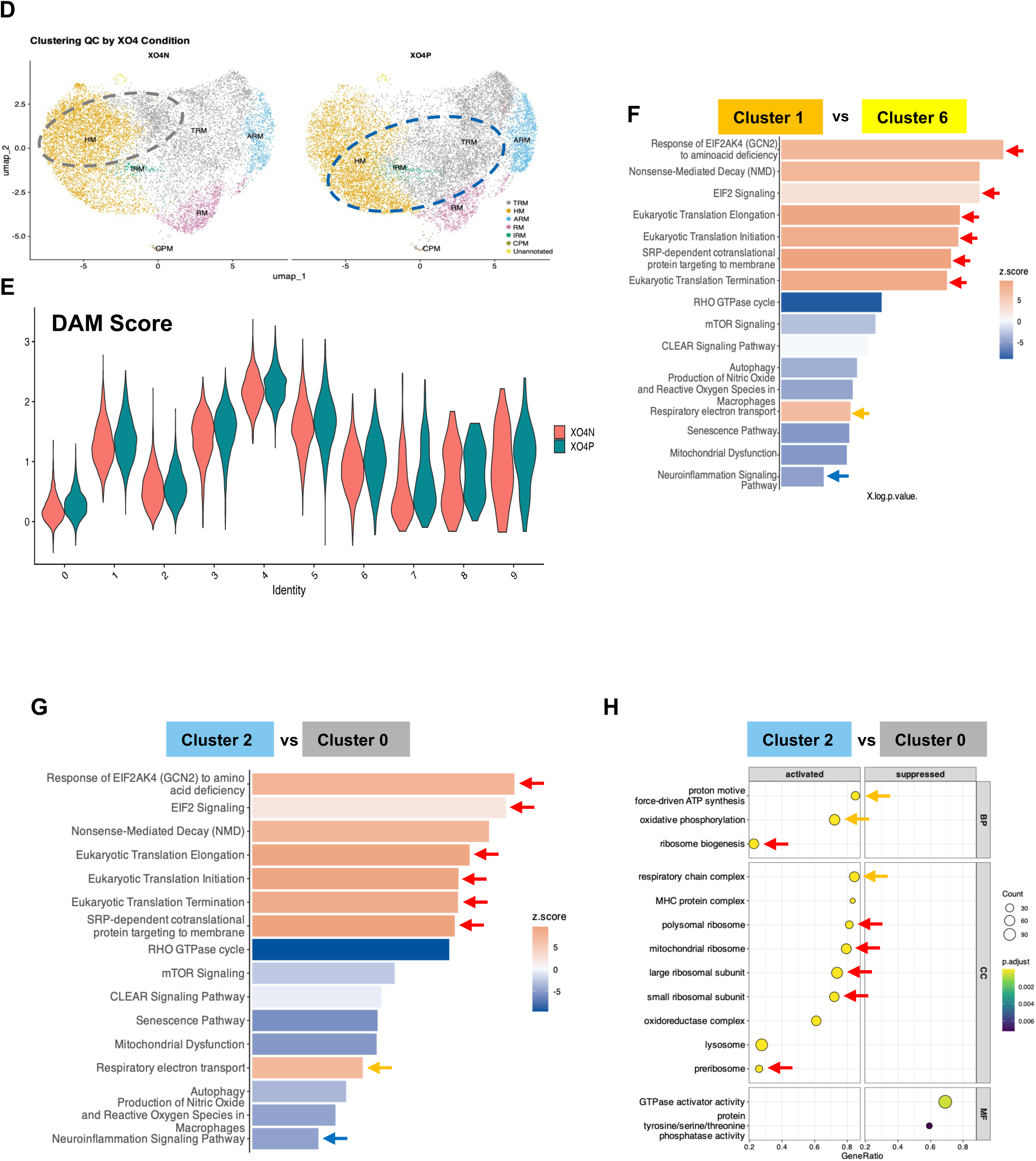
**(A)** Uniform Manifold Approximation and Projection (UMAP) of scRNA-seq data (resolution 0.2) identifies six distinct microglial clusters: Homeostatic Microglia (HM), Activated Response Microglia (ARM), Transition Response Microglia (TRM), Interferon Response Microglia (IRM), Cycling/Proliferating Microglia (CPM) (Sala Frigerio et al., 2019), and Ribosomal Microglia (Mancuso et al., 2024), along with a small unannotated cluster. **(B)** Validation of cluster identity using established module scores. Feature plots show the projection of gene signatures from Sala Frigerio et al. (2019), confirming consistency between the current clustering and previously defined states. Color gradients represent the calculated module score per cell (Blue: Low; Red: High). **(C)** Distribution of the canonical Disease-Associated Microglia (DAM) signature. UMAP projections (left) and violin plots (right) illustrate the expression of the DAM signature (Chen & Colonna, 2021) across non-phagocytic X04-and phagocytic X04 conditions. The y-axis represents the module score; violin width indicates cell density. **(D)** Clustering QC by X04 condition. UMAP projections of the high-resolution subclusters (resolution 0.3) split by non-phagocytic (X04-) and phagocytic (X04+) conditions, showing that X04+ and X04-cells occupy distinct subclusters. **(E)** Violin plots displaying DAM module scores for each subcluster (resolution 0.3) within X04-and X04+ conditions. Note that DAM scores are comparable between phagocytic and non-phagocytic subclusters. **(F)** IPA of top-ranked canonical pathways comparing the phagocytic Cluster 1 versus the non-phagocytic Cluster 6. Red arrows highlight the induction of protein-synthesis and translation pathways; the blue arrow denotes suppressed neuroinflammation signaling. **(G)** IPA of top-ranked canonical pathways comparing the phagocytic-enriched Cluster 2 versus the non-phagocytic Cluster 0. Red arrows highlight the upregulation of Ribosome Biogenesis and Protein Synthesis; yellow arrows denote the activation of Mitochondrial Oxidative Phosphorylation. **(H)** GSEA dot plot of Biological Process (BP) terms specifically enriched in Cluster 2 versus Cluster 0. Color gradient represents adjusted P-values.

**Extended Figure 6.**
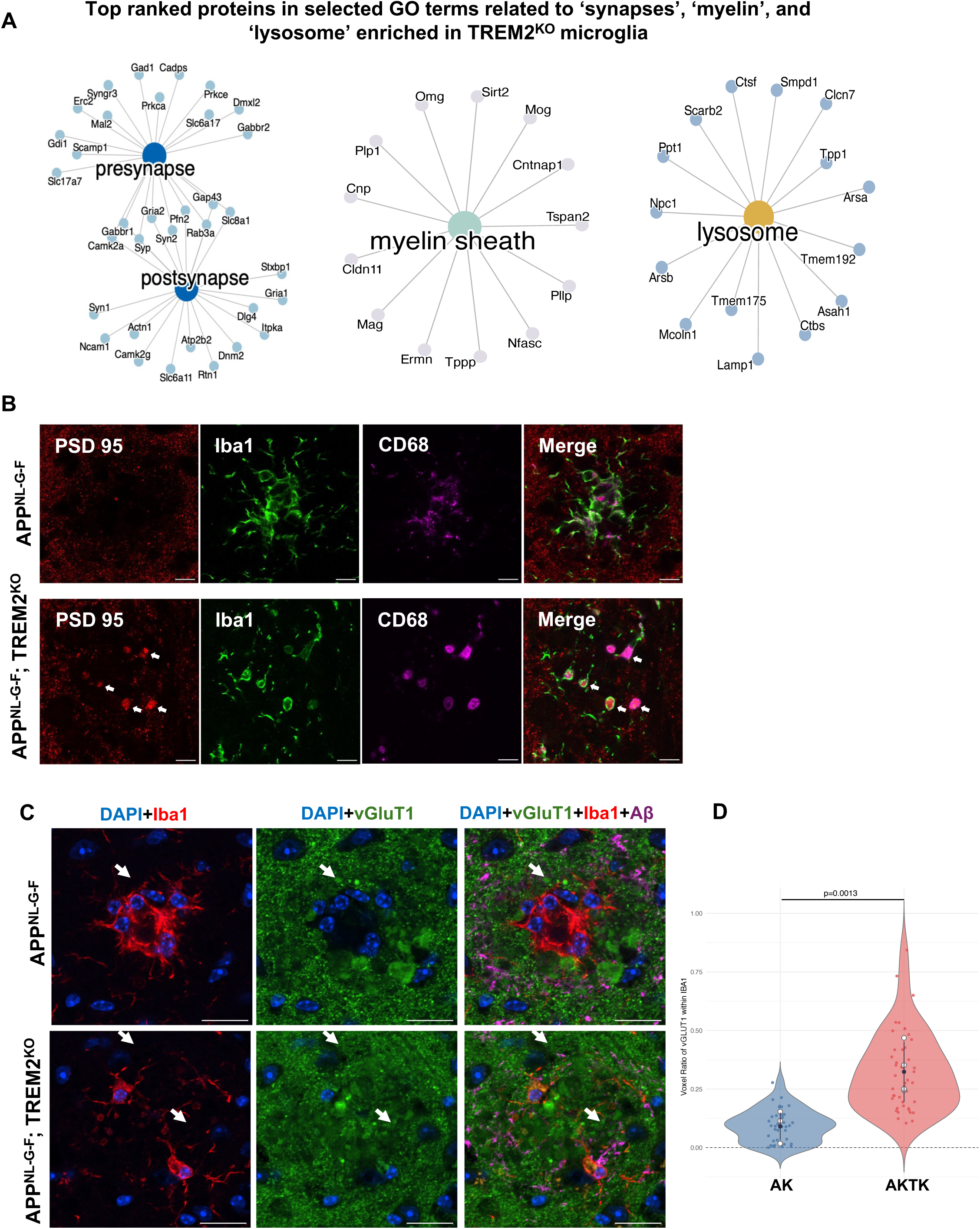
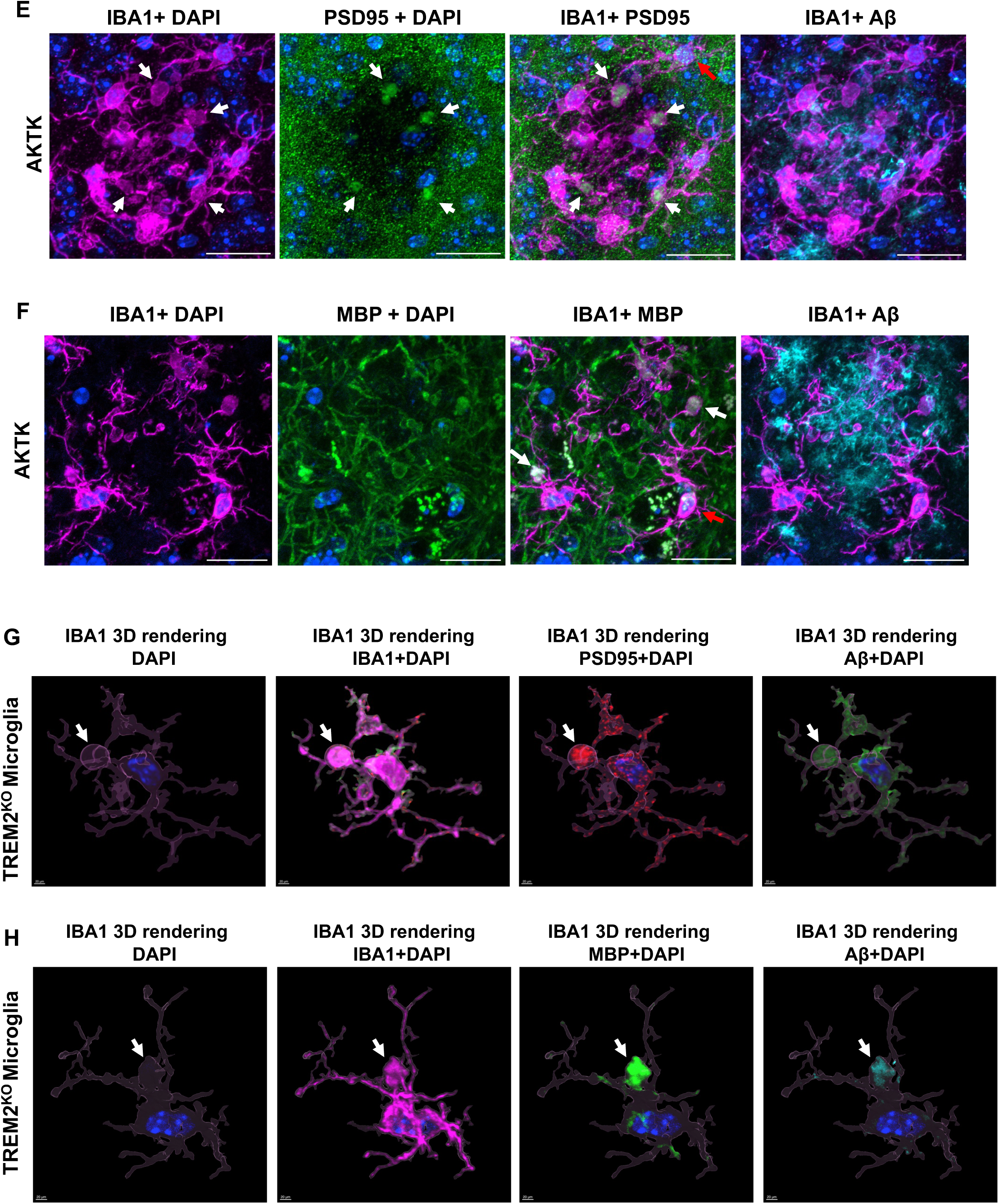
(A) Gene-concept network (cnet) plots displaying the top-ranked differentially abundant proteins associated with the Gene Ontology (GO) terms ‘Synapse’, ‘Myelin Sheath’, and ‘Lysosome’. Note that these cargo proteins are specifically enriched in *Trem2^KO^* microglia compared to *App^NL-G-F^*controls at 9 months of age, indicating a failure to degrade internalized material. **(B)** Representative confocal images of plaque-associated microglia in 9-month-old mice co-stained for PSD95 (postsynaptic marker), CD68 (lysosomal marker), and Iba1. White arrows highlight PSD95 puncta encapsulated within CD68+ phagolysosomes, indicating successful cargo internalization and trafficking to degradative compartments. Scale bar: 50 μm. **(C)** Representative confocal images of plaque-associated microglia in 9-month-old mice co-stained for vGluT1 (presynaptic marker), Iba1, and Aβ. White arrows highlight the pronounced intracellular accumulation of vGluT1 puncta within the soma of *Trem2^KO^*microglia. Scale bar: 50 μm. **(D)** Quantification of intracellular cargo accumulation (vGluT1 and others) in brain sections from 9-10 month-old *App^NL-G-F^* mice and *App^NL-G-F^;Trem2^KO^*mice. Colored points represent individual ROIs; solid black circles and error bars represent Estimated Marginal Means (EMMs) ± 95% CIs derived from a beta-binomial Generalized Linear Mixed Model (GLMM). (n = 5–6 mice per group). Pairwise comparisons were performed using the Delta method. Significance levels: *P < 0.05, **P < 0.01, ***P < 0.001, ****P < 0.0001. **(E)** Representative confocal images (MIP, 5 μm) of plaque-associated microglia co-stained for PSD95 (postsynaptic marker), Iba1, and Aβ. White arrows indicate large (>2 μm), process-associated cargo inclusions filled with PSD95 puncta emerging from Trem2^KO^ microglia. Red arrow indicates the cell used for the 3D surface rendering in Figure 6E. Scale bar: 20 μm. **(F)** Representative confocal images (MIP, 5 μm) co-stained for Myelin Basic Protein (MBP), Iba1, and Aβ. White arrows indicate MBP-filled cargo inclusions in Trem2^KO^ microglia. Red arrow indicates the cell used for the 3D surface rendering in Figure 6H. Scale bar: 20 μm. **(G-H)** Additional 3D surface renderings (Imaris) of plaque-associated microglia in *App^NL-G-F^;Trem2^KO^* mice, confirming the budding-like morphology of the cargo inclusions containing PSD95 (red, G) or MBP (green, H). The microglial membrane is rendered in magenta (Iba1).

**Extended Figure 7.**
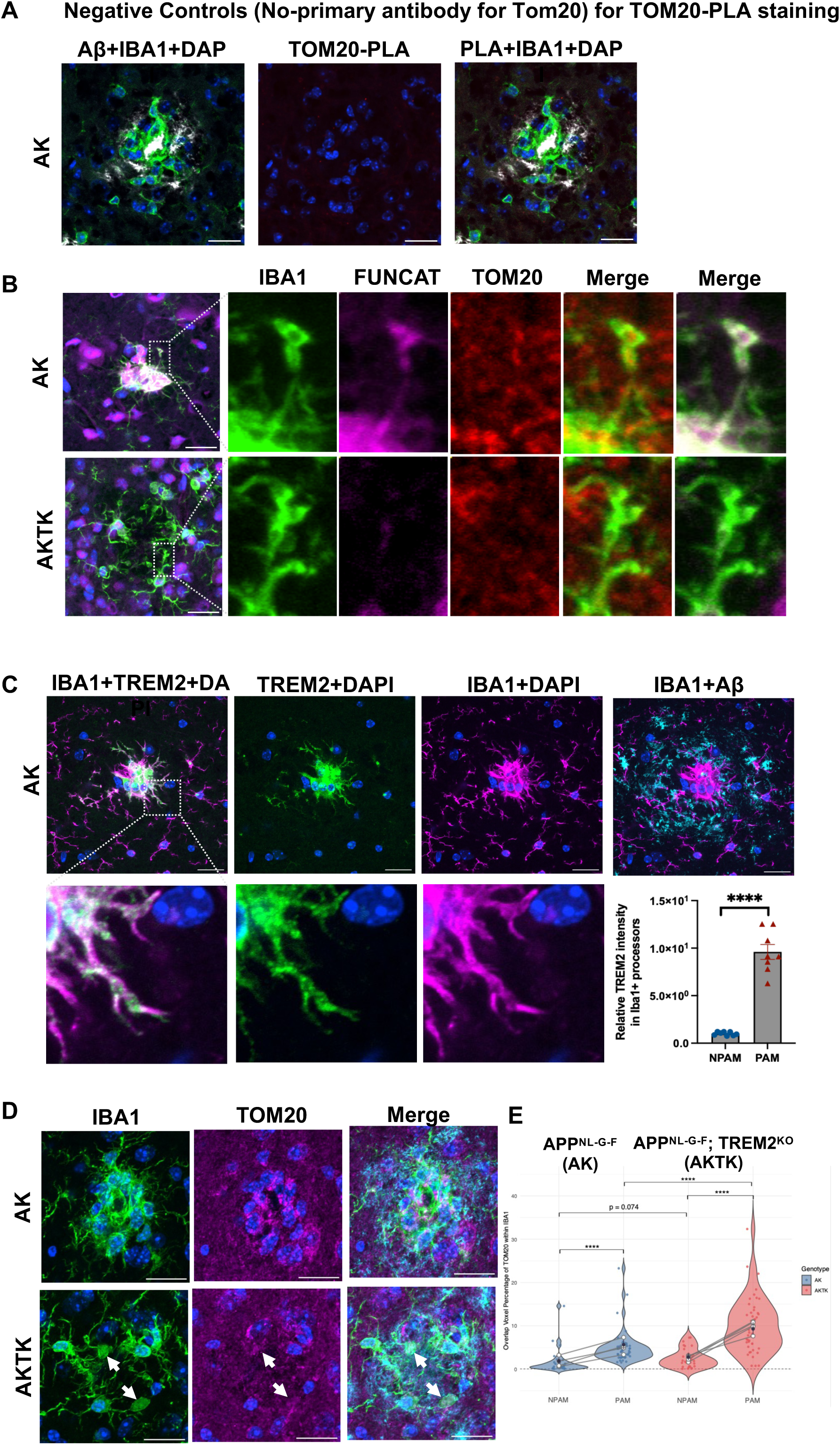
**(A)** Negative control for the TOM20 proximity ligation assay (PLA). Representative confocal images of plaque-associated microglia processed in parallel with the TOM20 primary antibody omitted; the absence of PLA signal confirms the specificity of the TOM20-PLA reaction. Scale bar: 20 μm. **(B)** Representative confocal images (MIP, 4 μm z-stack) of plaque-associated microglia in 9-10 month-old *App^NL-G-F^* mice and *App^NL-G-F^;Trem2^KO^* mice, co-stained for TOM20 (mitochondria), Iba1, and FUNCAT (nascent protein synthesis). Scale bar: 20 μm. **(C)** Representative confocal images (MIP, 4 μm z-stack) co-stained for TREM2, Iba1, and Aβ in both genotypes. White boxes highlight the specific induction of TREM2 protein within the processes of *App^NL-G-F^*microglia, spatially coinciding with the sites of local mitochondrial biogenesis shown in Fig. 7E. Scale bar: 20 μm. **(D-E)** Representative images and quantification of mitochondrial mass accumulation relative to plaque proximity. Violin plots display the distribution of TOM20 voxel overlap within Iba1+ ROIs, classified as Non-Plaque-Associated (NPAM) or Plaque-Associated (PAM). Colored points represent individual ROIs; solid circles and error bars represent Estimated Marginal Means (EMMs) ± 95% CIs derived from a beta-binomial Generalized Linear Mixed Model (GLMM). Lines connect paired regions within biological replicates (n = 5-6 mice). Pairwise comparisons were performed using the Delta method. ***P < 0.001, ****P < 0.0001.

**Extended Figure 8.**
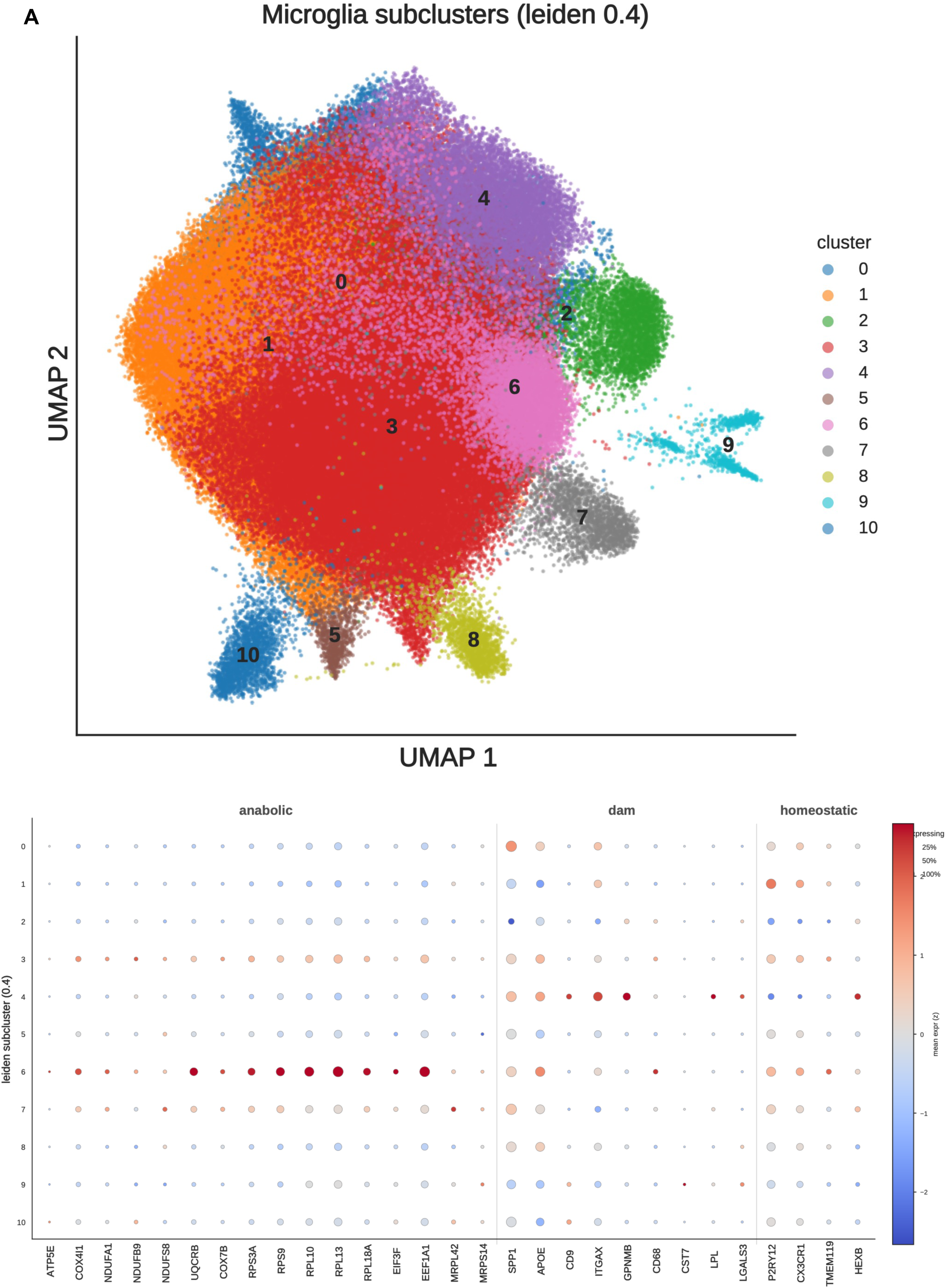
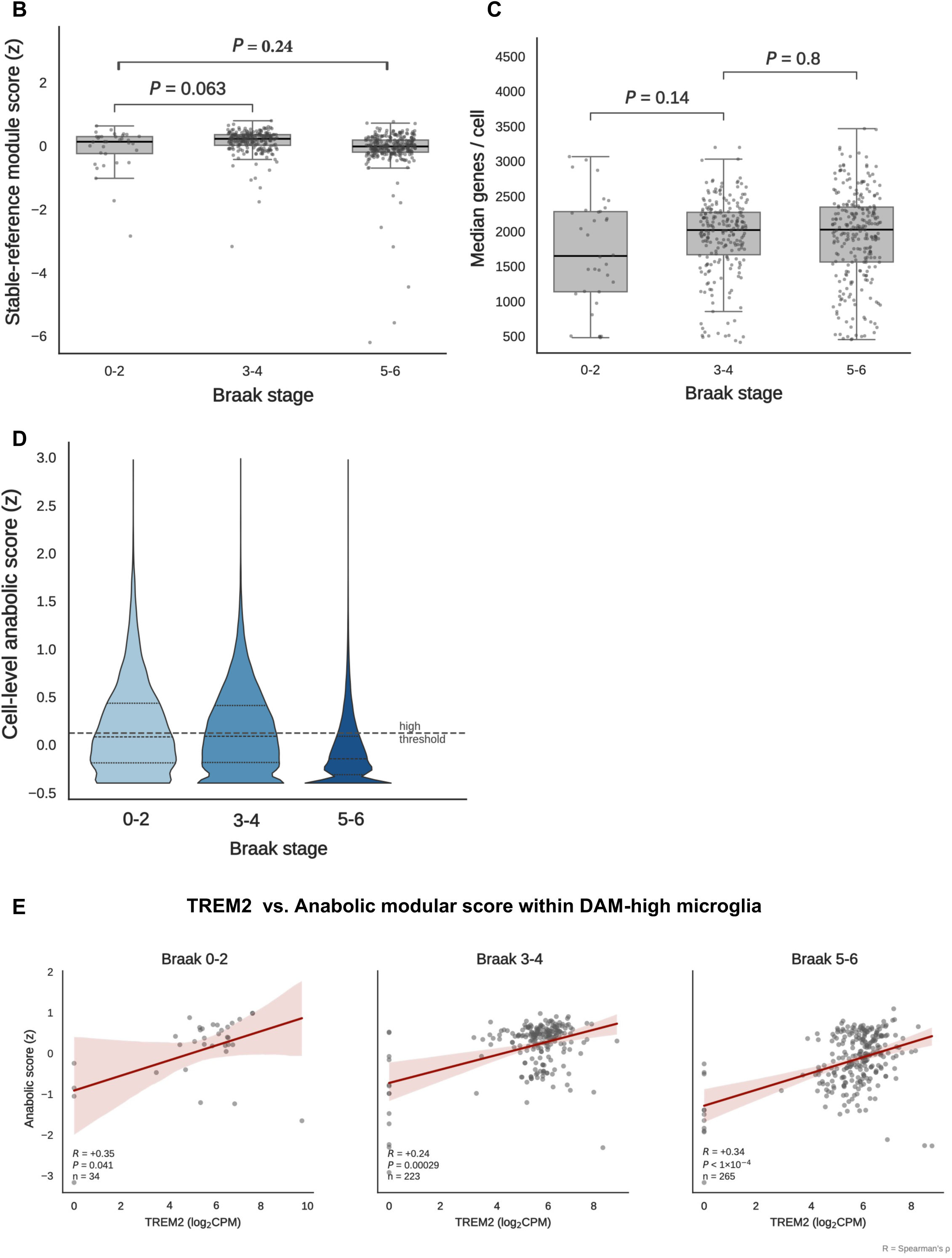
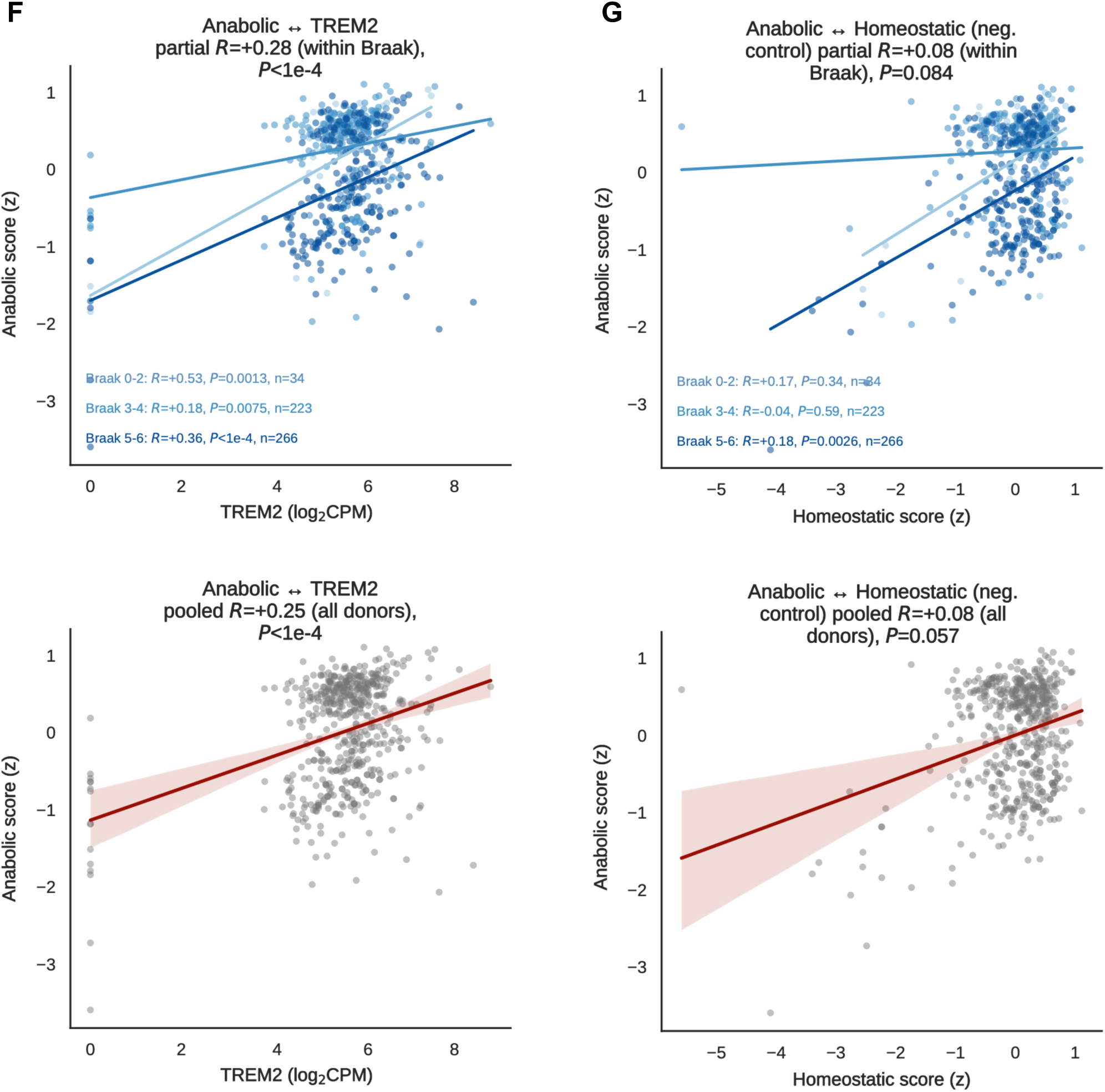
**(A)** Unsupervised re-clustering of human microglia (Leiden resolution 0.4) and module-gene dot plot, identifying a ribosomal/anabolic-high subcluster distinct from a DAM-marker-high subcluster. (**B**) Stable-reference control module score across Braak tiers (n.s.), based on nine widely used stable reference genes from the qPCR/RNA-seq normalization literature: ACTB, TBP, POLR2A, PPIA, HPRT1, YWHAZ, PUM1, CASC3, ELF1. (**C**) Median genes detected per nucleus across Braak tiers (sequencing-depth control; n.s.). (**D**) per-cell anabolic distribution by Braak. The donor-level decline is mainly compositional (fewer high-anabolic cells) with a smaller per-cell component. (**E**) TREM2 versus anabolic score within DAM-high microglia, per Braak stage (Spearman R). (**F**) Donor-level coupling of the anabolic program with TREM2. Top row: within-Braak regression with the partial Spearman R (controlling for Braak stage). Bottom row: the same relationship pooled across all donors (single regression, pooled Spearman R). (**G**) Donor-level coupling of the anabolic program with the homeostatic module (negative control). Top row: within-Braak regression with the partial Spearman R (controlling for Braak stage). Bottom row: the same relationship pooled across all donors (single regression, pooled Spearman R).

